# Pareto fronts reveal constraints on the evolution of ecological niches

**DOI:** 10.64898/2026.03.20.713179

**Authors:** Jason R. Laurich, Anita Narwani, Joey R. Bernhardt

**Affiliations:** Department of Integrative Biology, University of Guelph, Guelph, ON, Canada; Department of Aquatic Ecology, Swiss Federal Institute of Aquatic Science and Technology (Eawag), Dübendorf, Switzerland

**Keywords:** Evolutionary constraint, trade-offs, ecological limits, phytoplankton, global change

## Abstract

Trade-offs are central to biodiversity because they prevent the emergence of dominant phenotypes by limiting the simultaneous optimization of multiple fitness components. Yet trade-offs are often difficult to detect empirically when variation in overall performance produces positive correlations that mask underlying constraints. Here we use Pareto fronts—boundaries that capture optimal trade-off solutions—to test for evolutionary constraints on niche-determining traits in phytoplankton, including low-resource tolerance, thermal breadth, salt tolerance, and population growth rates. Using experimentally evolved *Chlamydomonas reinhardtii* populations subjected to nutrient and salt stress, we detected widespread Pareto fronts limiting the joint optimization of growth rate and niche-determining traits, thereby restricting the emergence of multivariate stress tolerance. Importantly, Pareto fronts revealed trade-offs even when underlying trait correlations were positive. We found that the structure of trait covariation behind Pareto fronts strongly predicted evolutionary outcomes: populations moved toward Pareto-optimal phenotypes primarily when trait correlations were neutral or positive, whereas negative trait correlations were associated with limited evolutionary optimization. Extending this framework across phytoplankton diversity, we compiled niche-determining traits for 189 phytoplankton species. At a macroevolutionary scale, we detected significant Pareto fronts constraining the evolution of niche-determining traits in phytoplankton. These fronts, however, did not always recapitulate the structure of trade-offs evident among *C. reinhardtii* populations, suggesting that forces that dictate microevolutionary outcomes, such as genetic correlations, may be resolved across macroevolutionary time. Together, our results highlight that evolutionary trajectories may differ across scales, but that fundamental limits on multivariate trait optimization persist across phytoplankton.

**Significance Statement:** Trade-offs among biological traits are central to evolutionary theory but often prove difficult to detect empirically. Here, we apply Pareto fronts—a framework borrowed from economics and engineering—to detect and reveal trade-offs among key niche-determining traits in phytoplankton. By combining experimental evolution in the laboratory with a synthesis of ecological traits across 189 species, we demonstrate widespread limits on the simultaneous optimization of growth rate, low-resource tolerance, salt tolerance, and thermal breadth. Importantly, Pareto fronts reveal trade-offs even when conventional correlation-based approaches fail, uncovering evolutionary constraints that remain hidden in trait correlations. These results show that trade-offs shape phenotypic variation across both micro- and macroevolutionary scales and impose fundamental limits on phytoplankton responses to multiple environmental stressors.

## Introduction

Since Darwin first remarked on his “tangled bank” of biodiversity (1), evolutionary ecologists have grappled with a central problem: identifying the forces that shape—and constrain—the patterns of variation that constitute Earth’s biodiversity (2). Among the most important of these forces are fundamental trade-offs between fitness-determining traits (1, 3–5). By preventing the simultaneous optimization of multiple traits, trade-offs have long been viewed as primary obstacles to the emergence of “Darwinian demons”—hypothetical genotypes capable of maximizing all aspects of fitness (6, 7). Variation in how organisms navigate these trade-offs is widely invoked to explain ecological specialization, niche differentiation, and the coexistence of competing species (8–11).

Beyond shaping patterns of biological variation, trade-offs are central to predicting evolutionary responses to environmental change. Natural selection acts on organisms through inherently multivariate selective landscapes, operating on interacting traits that together determine fitness (12). Consequently, evolutionary responses depend not only on the direction and strength of selection but also on the structure of trade-offs that limit the simultaneous optimization of multiple traits (13). The consequences of such trade-offs may be especially acute under anthropogenic change, as human activities increasingly expose organisms to co-occurring stressors including warming, salinization, pollution, and altered nutrient regimes (14–17).

Primary producers such as phytoplankton, which underpin aquatic food webs and drive global biogeochemical cycles (18, 19), face complex multivariate selection acting on key niche-determining traits. Traits including growth rate, minimum resource requirements (*R*^*^), thermal breadth, and salt tolerance play central roles in defining phytoplankton niches and species persistence under environmental change and are hypothesized to be linked by trade-offs (11). Classic examples include gleaner-opportunist (20–23), tolerance–growth (24, 25), and generalist–specialist (26) trade-offs within individual niche axes. Trade-offs may also arise across niche axes—for example between performance under low-light and low-nutrient conditions—depending on whether traits share physiological pathways or compete for limited energetic or cellular resources (27-30). Where present, such trade-offs should exert powerful constraints on evolutionary responses to environmental change. Yet despite their central role in ecological and evolutionary theory, the extent to which trade-offs shape phenotypic variation in natural systems remains uncertain, and clear empirical evidence for them is often elusive (3, 5, 31).

Trade-offs are frequently obscured in empirical data, where traits presumed to trade off often exhibit positive correlations instead (3, 5, 31, 32). This pattern can arise when individuals vary widely in overall performance due to physiological, genetic, and environmental differences (33) and high performers excel across multiple tasks simultaneously, masking underlying constraints (Figure 1AB). Nevertheless, theory predicts that no genotype can simultaneously maximize all aspects of fitness (6, 34, 35). Trade-offs can therefore become apparent among relatively optimized phenotypes (3, 36-38), where constraints emerge as Pareto fronts—the outer boundary of achievable trait combinations beyond which one trait cannot improve without another declining. First formalized in economics (39) and adapted to solve multi-objective optimization problems in engineering (40, 41), Pareto fronts—applied to evolutionary ecology—characterize ceilings on natural selection’s capacity to jointly optimize biological tasks and traits (35, 42), rendering regions of phenotypic space effectively inaccessible (Figure 1C). Given that Pareto fronts represent the set of optimally efficient solutions to a trade-off problem, rather than the average relationship across all phenotypes, they can reveal trade-offs even where the overall correlation is positive or neutral (Figure 1AB) —precisely the cases in which traditional correlational approaches often fail (2, 3, 33).

**Figure 1.**
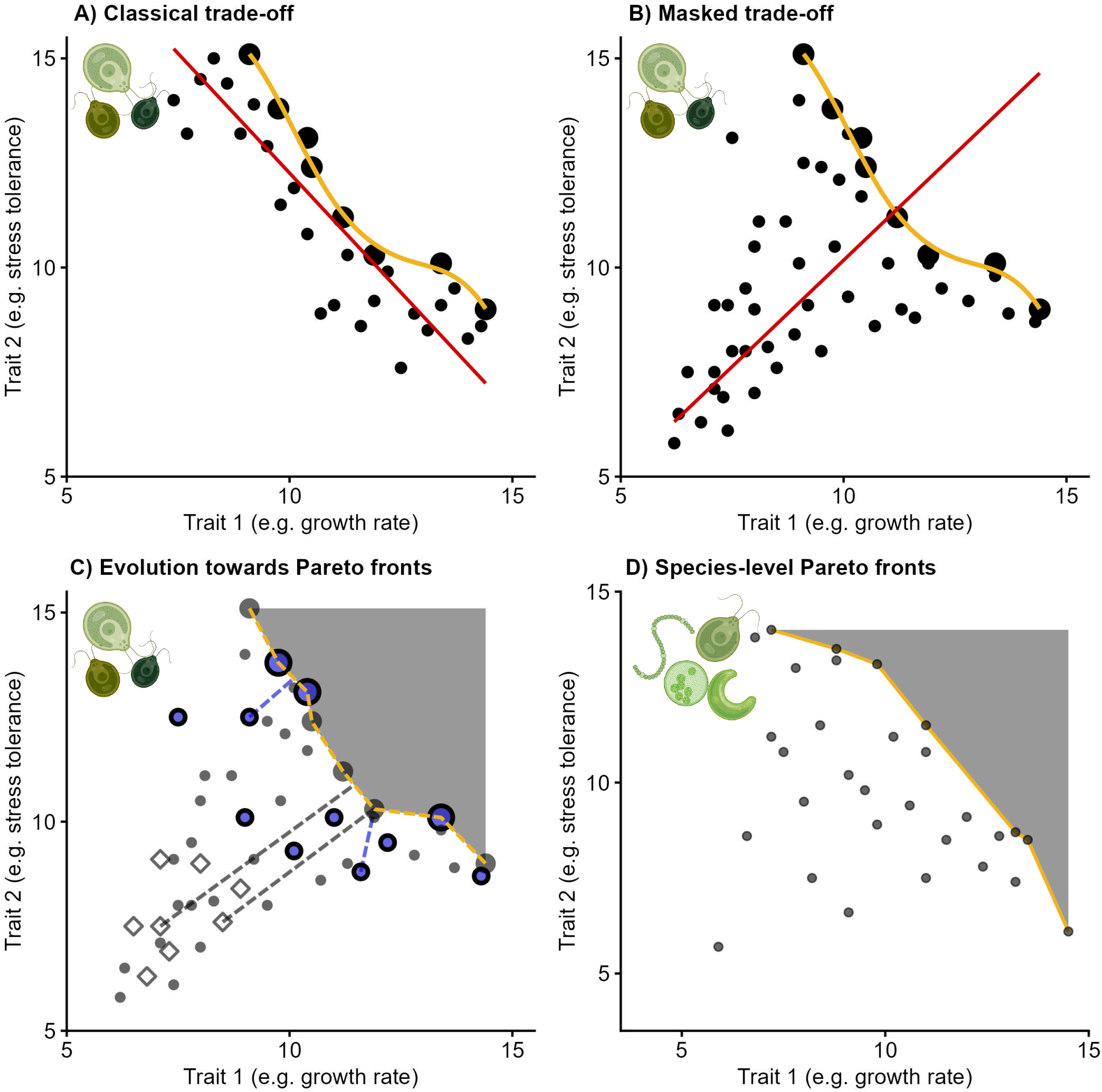
Representation of trade-off configurations and detection methods using hypothetical data. Phytoplankton icons depict the relevant level of biological organization: within a species (A-C) or across multiple species (D). (A) Classical trade-offs manifest as direct negative correlations between competing traits (red line). In such cases, Pareto fronts—the set of trait combinations that optimally balance conflicting demands (yellow)—are not required to detect trade-offs. (B) When variation in individual quality (33), inconsistent selection, or multivariate trade-offs invisible in two-dimensional plots obscure a trade-off, raw correlations (red line) fail to recover the trade-off. Pareto fronts (yellow) can instead identify the distribution of optimal trait combinations that define the underlying trade-off surface. (C) Pareto fronts constrain evolutionary optimization by generating inaccessible regions of phenotypic space (dark grey). In addition to quantifying this inaccessible space, we tested whether experimental evolution moves populations towards Pareto-optimal solutions by calculating the Euclidean distance of each ancestral and descendant population from the Pareto front (see Methods). Hypothetical ancestral trait combinations are shown as open diamonds connected to the front via grey dashed lines; hypothetical descendant populations evolved under selection to optimize trait values (‘matching populations’) have moved towards the Pareto front and are shown as black-outlined blue circles connected to the front via blue lines. For visual clarity, these connecting lines are only shown for a subset of points. (D) Across species, Pareto fronts define the boundary of accessible trait space (yellow line), creating a region of optimized trait space that no species can occupy (grey region). In panel D, each point represents the average trait value of a species.

Populations under directional selection are expected to evolve towards Pareto fronts, optimizing trait values until they reach the boundary (Figure 1C). For genotypes located on or near a Pareto front, simultaneous improvement in multiple traits is expected to be exceedingly unlikely (42, 43). Not all genotypes, however, are expected to lie on a given front. Because a Pareto front is a two dimensional projection of higher-dimensional trade-offs, genotypes that appear sub-optimal for a given trait pair may in fact be optimized along additional axes of selection (35). These three-(or more)-dimensional trade-offs will manifest as constraint envelopes with multiple poles in bivariate space, each representing an archetype specialized for one task at the expense of others (10, 24, 27, 35). Phenotypes in the interior of these spaces may thus be generalists that balance investment across several traits (Figure 1B). Over longer timescales, sufficiently strong or novel selection pressures may occasionally shift the location of a Pareto front by favoring mutations that improve performance across multiple traits simultaneously, particularly when constraints imposed by these other aspects of organismal biology — such as cell size, defense, or additional niche axes in phytoplankton (23, 27, 29, 44) — are relaxed.

Across macroevolutionary timescales, selection acting on many traits across diverse lineages should expose fundamental trade-offs that remain unresolved (5). When selection strongly favours optimization along a particular pair of traits, species are predicted to sort along Pareto fronts, producing negative correlations near the upper bounds of performance (45–49). In contrast, broad patterns of specialization and divergence in ecological niches and life history strategies reflect the historical imprint of selection acting on many traits at once — divergence often driven by major innovations in life history and physiology (50), including multicellularity (51), coloniality (52), and nitrogen fixation (53) in phytoplankton, that open new ecological strategies and expand the achievable phenotypic space. Yet however much such innovations expand the range of viable strategies, the outer bounds of occupiable phenotypic space should still reflect hard limits on phenotypic optimization. Because genetic constraints are less relevant over long evolutionary timescales (54 –57), these macroevolutionary patterns are expected to reveal fundamental biophysical or physiological limits to trait optimization, rather than the transient genetic constraints and historical contingency that shape variation within species (3, 31).

In this study, we combined experimental evolution and comparative data synthesis to examine how Pareto fronts shape patterns of phenotypic variation and constraint across micro- and macroevolutionary scales in phytoplankton. Trade-offs inferred from observational or comparative data are inherently difficult to interpret (22, 23, 31), because correlations can reflect genuine biological constraints or simply adaptation to correlated environmental gradients along which species are distributed (e.g. the covariation between temperature and thermal heterogeneity across latitudes; 58–60). Experimental evolution circumvents this ambiguity by imposing controlled, independent selection on populations derived from common ancestors, thus isolating the genetic and physiological trade-offs that remain once environmental covariation is removed. Using 37 experimentally evolved populations of *Chlamydomonas reinhardtii*, we first tested (1) whether Pareto fronts constrain relationships between growth and key niche-determining traits within and across environmental gradients, and (2) under what conditions experimental evolution drives populations toward Pareto-optimal trait combinations. We then extended this framework to a synthesis of niche-determining traits across 189 phytoplankton species to test (3) whether the signatures of Pareto constraint observed in experimentally evolved *C. reinhardtii* populations are reflected in macroevolutionary patterns of trait variation. Throughout, we focus on trade-offs involving low-resource tolerance, salt tolerance, and thermal performance—traits central to defining phytoplankton niches under contemporary environmental change.

## Results

### Trade-offs between growth and niche-determining traits in *Chlamydomonas reinhardtii*

To investigate trade-offs between maximum growth rates and key niche-determining traits, we experimentally evolved five ancestral populations of *Chlamydomonas reinhardtii* under salt stress and light, nitrogen, and phosphorus limitation (Supporting Methods 2.i-ii). After approximately 285 generations, we assayed ancestral and descendant populations across gradients of light, nitrogen, phosphorus, salt, and temperature—though no populations evolved under thermal stress—to estimate maximum growth rates (*µ*_*max*_), low-resource tolerance, salt tolerance (*S*), and thermal breadth (*T*_*br*_) (Figure S1). We defined low-resource tolerance as 1/*R*^*\**^ (21), the inverse of the minimum resource level required for population growth (*R*^*\**^). In total, these assays comprised approximately 5,700 independent time series of population abundances across 6-10 levels of five distinct environmental gradients (light, nitrogen, phosphorus, salt, and temperature), which we used to estimate growth rates. We then fit roughly 630 Bayesian models that capture variation in population growth rates across these abiotic gradients. Each trait estimate used to investigate trade-offs among niche-determining traits thus represents a curve fit to 6-10 growth rates, each derived from its own population abundance time series.

We focused first on trade-offs between *C. reinhardtii* growth rates and niche-determining traits within individual niche axes. Here, classical ecological trade-offs such as gleaner-opportunist, generalist-specialist, and growth-tolerance trade-offs are expected to arise directly from physiological constraints that limit simultaneous optimization along a single dimension of environmental variation—for example when adaptations that improve growth under low resource availability reduce maximum growth rate when that resource is replete (20, 21, 24–26). To identify trade-offs among phenotypes approaching optimal trait combinations, we characterized putative Pareto fronts using a modified convex hull algorithm and fit shape-constrained additive models to Pareto-optimal phenotypes. We assessed evidence for the existence of these putative Pareto fronts by calculating the empty phenotypic space they render inaccessible (Figure 1C) and comparing our observations to randomized datasets using permutation tests. We also computed Pearson correlations (*r*) to assess the strength and direction of overall trait covariation—a traditional, correlational counterpart to our Pareto front framework.

Pareto front analyses revealed strong constraints on the joint optimization of *µ*_*max*_ and niche-determining traits across all environmental gradients we considered (Figure 2A-E). In each case, we found that phenotypic distributions were bounded by Pareto fronts, indicating trade-offs between maximum growth rate and niche-determining traits. The area of phenotypic space excluded by these fronts significantly exceeded randomized null expectations (42) for the relationships between *µ*_*max*_ and low-L tolerance (one-tailed permutation test, *P* < 0.001, Figure 2A), low-N tolerance (*P* < 0.001, Figure 2B), low-P tolerance (*P* < 0.001, Figure 2C), salt tolerance (*P* = 0.008, Figure 2D), and thermal breadth (*P* < 0.001, Figure 2E). Critically, we also found that Pearson correlations between *µ*_*max*_ and niche-determining traits were significantly negative within individual niche axes (Figures 2A-E, 5A top left quadrant), indicating that these trade-offs extend broadly across phenotypic space in *C. reinhardtii*, rather than occurring only among Pareto-optimal phenotypes.

**Figure 2.**
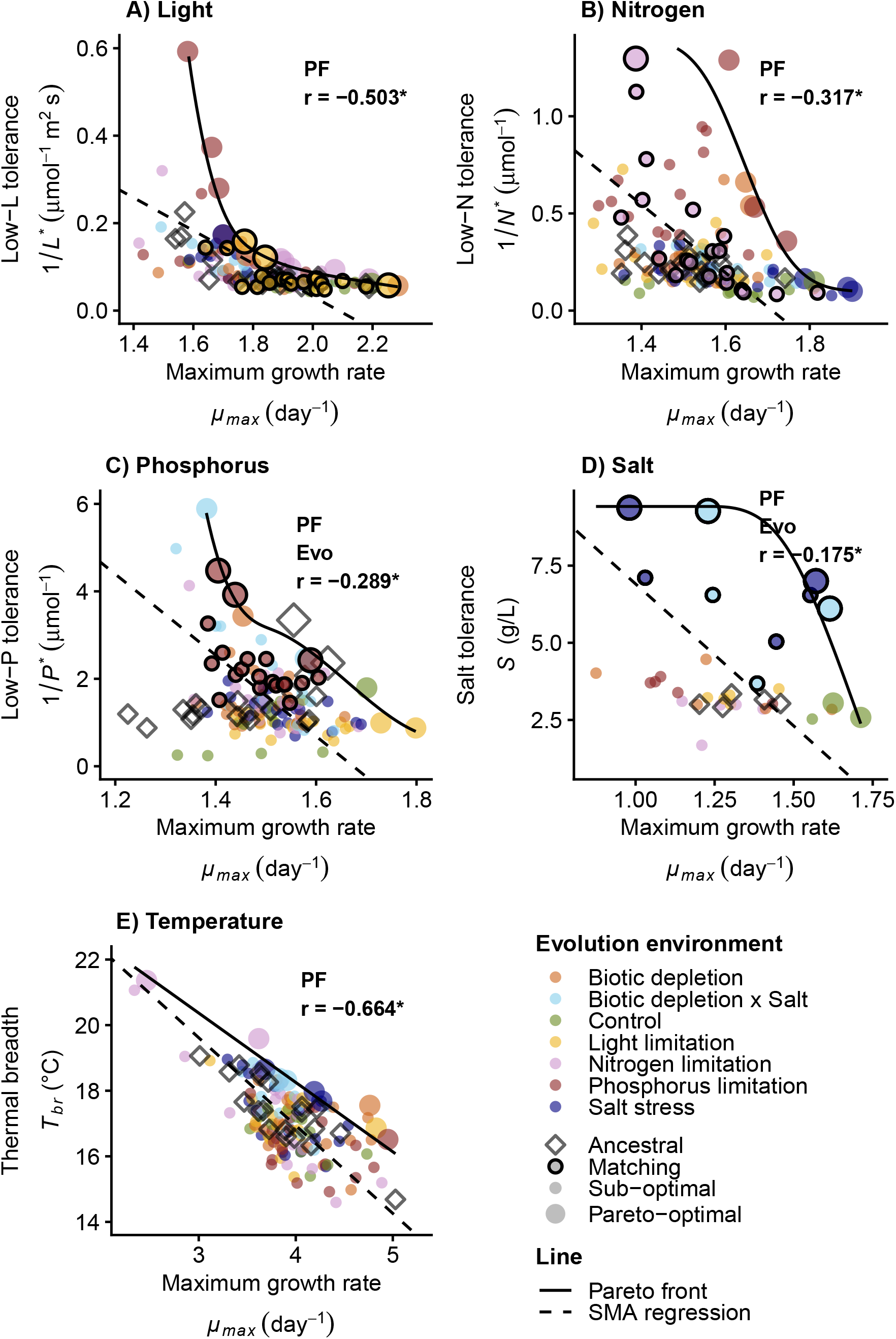
Trade-offs between maximum specific growth rates (*μ*_*max*_) and niche-determining traits in experimental *Chlamydomonas reinhardtii* populations, plotted using all trait data. (A-C) Trade-offs between low-resource tolerance (1/*R*^*\**^) and *μ*_*max*_ for light, nitrogen, and phosphorus. (D) Trade-off between salt tolerance (S) and *μ*_*max*_. (E) Trade-off between thermal breadth (*T*_*br*_) and *μ*_*max*_. Trait data for populations that evolved under the panel-relevant selective environment (e.g. under light limitation in A) are shown as filled circles outlined in black (“Matching”); other experimentally evolved populations are filled circles without outlines. Each evolutionary treatment comprised five populations, except the control (N = 3) and biotic depletion × salt (N = 4); five ancestral populations are shown as open diamonds. Each point represents trait data from one of four replicate assays per population (salt assays were not replicated). Smoothed black lines—representing putative Pareto fronts—were fit to the set of optimal trait combinations (identified via a convex hull algorithm) using smoothed, monotonically decreasing curves. Pareto-optimal trait combinations are shown as larger circles and diamonds, while the smaller points refer to “sub-optimal” trait combinations. Dashed lines are standardized major axis (SMA) regressions, plotted only where the Pearson correlation between traits was significant (r reported in each panel; asterisk denotes significance at *α* = 0.05). All panels had statistically significant Pareto fronts (which define larger inaccessible spaces than null, randomized simulations) and are plotted in black. Text labels report the correlation (*r*) and significant results (*α* = 0.05; PF: a significant Pareto front constrains phenotypic variation; Evo: experimental evolution has moved matching populations significantly closer to the Pareto front).

### Pareto fronts reveal trade-offs among niche-determining traits in *Chlamydomonas reinhardtii*

Whereas trade-offs between growth and niche-determining traits within a single niche axis are expected to arise from physiological constraints, relationships among traits governing performance across multiple niche axes (e.g. salt and temperature) may be shaped by a broader and less predictable set of forces including genetic, physiological, and resource-allocation trade-offs. These forces need not act in concert: antagonistic pleiotropy or competition for a shared resource budget can generate negative correlations (12, 13, 23, 27), synergistic pleiotropy can improve multiple traits simultaneously, and modular traits can evolve independently (21, 43). Cross-axis relationships therefore carry no consistent expectation of sign—making the Pareto front approach, which detects constraints regardless of the underlying correlation, especially informative.

Pareto front analyses revealed several constraints on the simultaneous optimization of niche-determining traits across niche axes (Figure 3). Significant Pareto fronts bounded the thermal breadth (*T*_*br*_)-low-L tolerance (one-tailed permutation test, *P* < 0.001, Figure 3D), *T*_*br*_,-salt tolerance (*P* < 0.001, Figure 3J), low-N tolerance-salt tolerance (*P* < 0.001, Figure 3F), and low-P-low-L tolerance (*P* = 0.012, Figure 3B) relationships. We detected additional weaker fronts between low-p tolerance and low-N tolerance (*P* = 0.047, Figure 3E), and salt tolerance (*P* = 0.043, Figure 3H). These latter fronts sat near the boundary of statistical significance (*α* = 0.05). Because Pareto fronts describe only relatively high-performing trait combinations, we tested them using only the most optimized third of observations—those furthest, by scaled Euclidean distance, from the minimum (x, y) values. The significance of these weaker front–but not the stronger fronts discussed above—depended on our choice of threshold (Figure S2, Table S1; see Methods).

**Figure 3.**
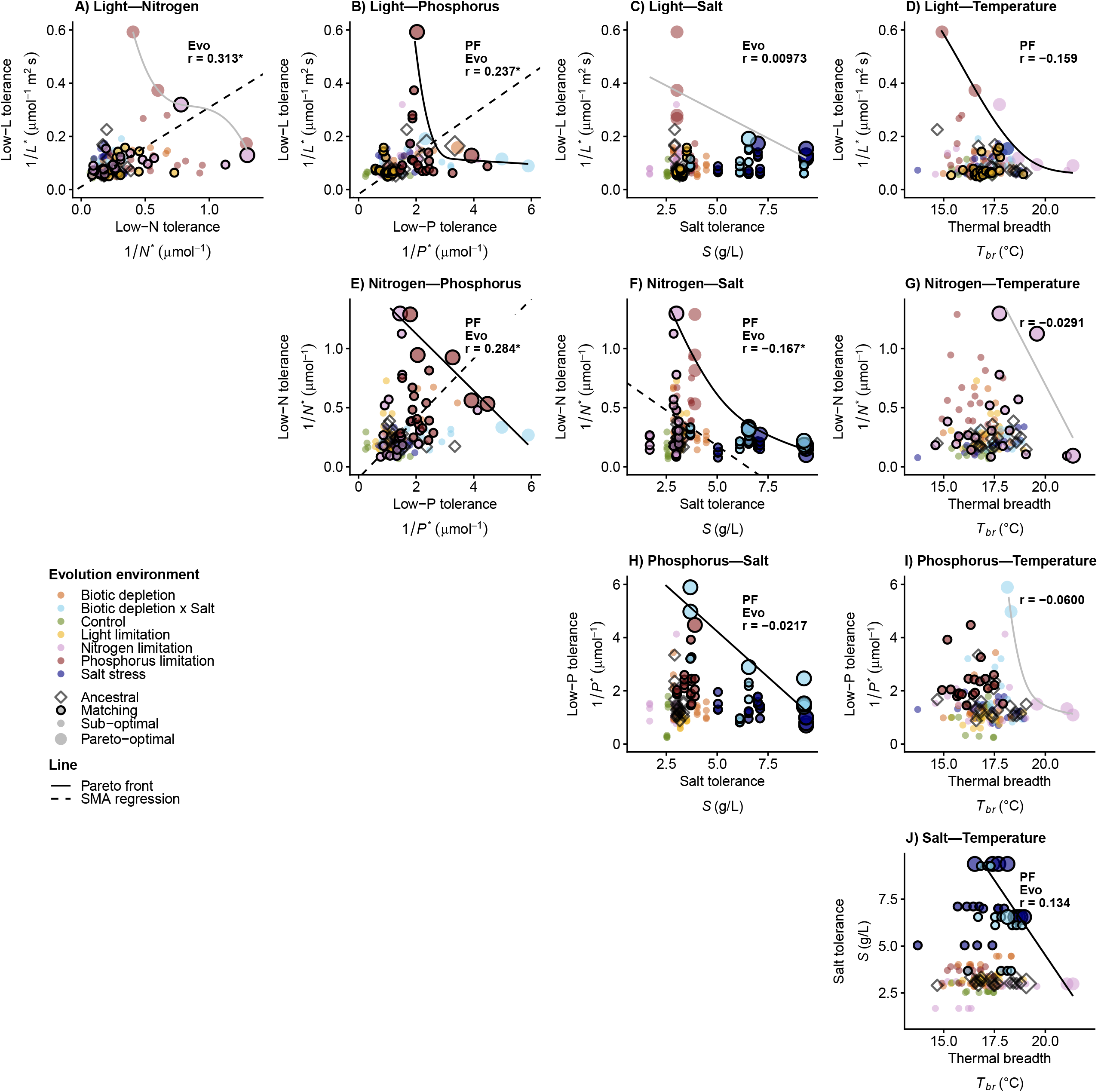
(A-J) Trade-offs between low-resource tolerance (1/*R*^*\**^) for light, nitrogen, and phosphorus, thermal breadth (*T*_*br*_) and salt tolerance (S) in experimentally evolved *Chlamydomonas reinhardtii* populations. Niche-determining traits are plotted in a single row and/or column. Trait data for populations that evolved under the panel-relevant selective environment (e.g. under light OR nitrogen limitation in A) are shown as filled circles outlined in black; other experimentally evolved populations are filled circles without outlines. Each evolutionary treatment comprised five populations, except the control (N = 3) and biotic depletion × salt (N = 4); five ancestral populations are shown as open diamonds. Each point represents trait data from one of four replicate assays per population (except for salt assays, which were not replicated). Smoothed solid lines—representing putative Pareto fronts—were fit to the set of optimal trait combinations (identified via a convex hull algorithm) using smoothed, monotonically decreasing curves. Pareto-optimal trait combinations are shown as larger circles and diamonds, while the smaller points refer to “sub-optimal” trait combinations. Dashed lines are standardized major axis (SMA) regressions, plotted only where the Pearson correlation between traits was significant (*r* reported in each panel; asterisk denotes significance at *α* = 0.05). Significant Pareto fronts (which define larger inaccessible spaces than null, randomized simulations) are plotted in solid black lines; non-significant Pareto fronts are shown in solid grey lines. Text labels report the correlation (*r*) and significant results (*α* = 0.05; PF: a significant Pareto front constrains phenotypic variation; Evo: experimental evolution has moved at least one matching population significantly closer to the Pareto front). All data are plotted.

Most strikingly, several fronts emerged despite positive or neutral correlations in the underlying trait data (e.g. the low-P-low-L tolerance, low-P-low-N tolerance, and salt tolerance-thermal breadth relationships; Figures 3BEJ, 5A, Table S2). In these cases, correlational approaches failed to detect trade-offs, yet Pareto fronts revealed constraints on joint trait optimization—demonstrating the utility of this framework in detecting multivariate constraints in the face of positive or neutral trait covariance.

### Evolutionary shifts towards Pareto-optimal trait combinations

Detecting Pareto fronts reveals constraints on multivariate trait optimization but does not describe whether evolution drives populations towards these boundaries. We therefore asked whether experimental evolution moved populations towards Pareto fronts—specifically, whether populations that evolved under a given selective environment (‘matching populations’) became better optimized for the traits governing performance in that environment. For each front identified within (Figure 2) and across (Figure 3) niche axes, we measured the distance of each population from the putative Pareto front in standardized (z-scored) trait space and used mixed-effects models to test whether matching evolved populations lay significantly closer to the Pareto front than their ancestors (Figures 1C, S3).

Experimental evolution produced clear, condition-specific shifts toward some Pareto fronts. Populations that evolved under salt stress were significantly closer to the *µ*_*max*_-salt tolerance front than their ancestors (linear mixed-effects model, one-tailed contrasts, *P* < 0.001, Figure 2D), and populations that evolved under P-limitation also lay closer to the *µ*_*max*_ *-1/P*^*^ front than their ancestors (*P* < 0.001, Figure 2C). We detected no comparable evidence of optimization for the other within-axis Pareto fronts (Figure 2). Across niche axes, populations that evolved under matching conditions lay significantly closer to their Pareto fronts than their ancestors for most (7/10) relationships (*P* < 0.05, Figure 3). Strikingly, six of these seven relationships involved either salt tolerance or low-P tolerance—the two most evolutionarily responsive traits in our experiment (Tables S3-4, Figure S4) — the sole exception being low-L-low-N tolerance.

The extent to which evolution produced joint trait optimization was strongly influenced by the structure of trait covariance behind a Pareto front (Figure 5B). Where underlying trait correlations were positive, populations readily optimized multiple traits together through correlated evolution (e.g. low-P with low-L and low-N tolerance, Figure 3BE). Populations were also often able to optimize performance across multiple niche axes when trait correlations were near-zero (e.g. Figures 3CHJ, 5B). Significant negative correlations, by contrast, were generally associated with little movement toward Pareto fronts (Figures 2, 5B). The clearest exceptions were the two most evolutionarily labile traits: salt tolerance — which increased strongly in salt-adapted populations regardless of its correlations with other traits, a signature of modular evolution — and low-P tolerance, both of which drove optimization even against negative covariance (Figures 2, 3, 5B, Table S3). Across all analyses, evolutionary environment explained variation in niche-determining traits far better than ancestry (redundancy analysis, Figure S5; mixed-effects models, Table S4), reflecting adaptive responses to selection rather than differences in ancestral genetic background.

### Pareto fronts reveal broad trade-offs across phytoplankton niche axes

To evaluate whether Pareto constraints observed in our experimental evolution extend across macroevolutionary scales, we compiled estimates of growth rates and niche-determining traits across light, nitrogen, phosphorus, and temperature gradients for a broad range of phytoplankton taxa. After fitting thermal performance and Monod curves and screening model fits for quality (Supporting Methods 2.xiii), our dataset comprised 499 individual models spanning 189 species. To evaluate constraint in this broader, cross-species phenotypic space, we applied Pareto front analyses to bivariate relationships using mean trait values for each species.

Across species, we detected strong evidence of Pareto fronts that limit the simultaneous optimization of multiple niche-determining traits. Within single niche axes, we found that Pareto fronts constrained the *μ*_*max*_ low-L tolerance (one-tailed permutation test, *P* = 0.002, Figure 4A), *μ*_*max*_-low-P tolerance (*P* < 0.001, Figure 4H), and *μ*_*max*_-thermal breadth (*P* = 0.022, Figure 4J) relationships. We also found significant evidence of Pareto fronts that generate empty phenotypic space across phytoplankton niche axes, in the low-L-low-N tolerance (*P* = 0.002, Figure 4B), low-L-low-P tolerance (*P* = 0.010, Figure 4C), low-L tolerance-thermal breadth (*P* = 0.005, Figure 4D), and low-N tolerance-thermal breadth (*P* < 0.001, Figure 4G) relationships. In many cases, these cross-gradient constraints emerged only after excluding a single high-performing outlier species (>90th quantile on both niche axes), whose inclusion would otherwise obscure the broader pattern of constraint (Figure 4).

**Figure 4.**
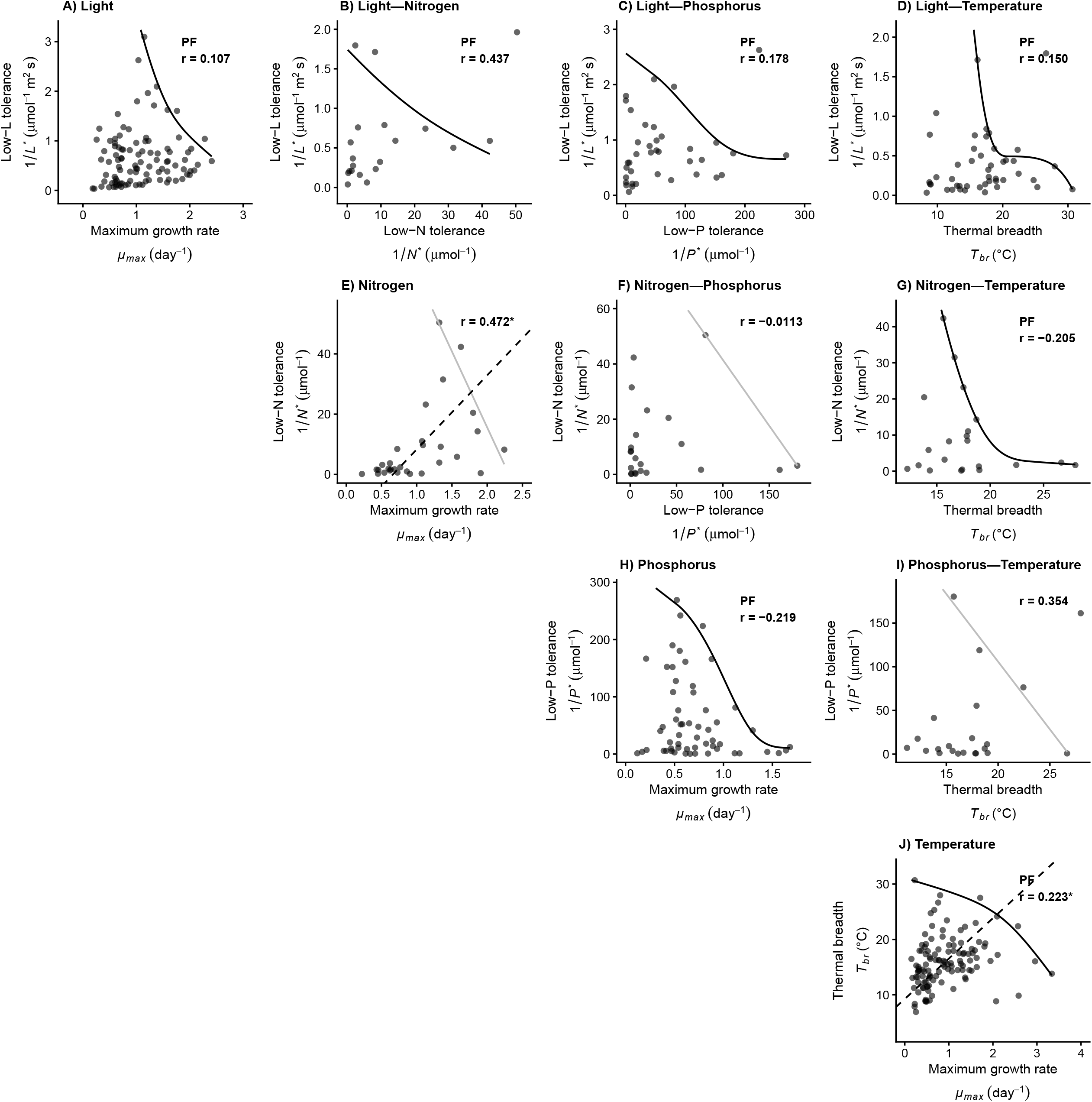
Trade-offs between maximum specific growth rates (*μ*_*max*_), low-resource tolerance (1/*R*^*\**^) for light, nitrogen, and phosphorus, and thermal breadth (*T*_*br*_) across 189 phytoplankton species. All data points represent the mean trait values of a single phytoplankton species; apparent variation in point shading reflects the plotting of overlapping values. Solid lines represent bivariate Pareto fronts constraining trait co-optimization using smoothed, monotonically decreasing curves. Dashed lines are standardized major axis (SMA) regressions, plotted only where the Pearson correlation between traits was significant (*r* reported in each panel; asterisk denotes significance at *α* = 0.05). Text labels report the correlation (*r*) and significant results (*α* = 0.05; PF: a significant Pareto front constrains phenotypic variation). Significant Pareto fronts (which define larger inaccessible spaces than null, randomized simulations) are shown in solid black lines; non-significant fronts are light grey.

**Figure 5.**
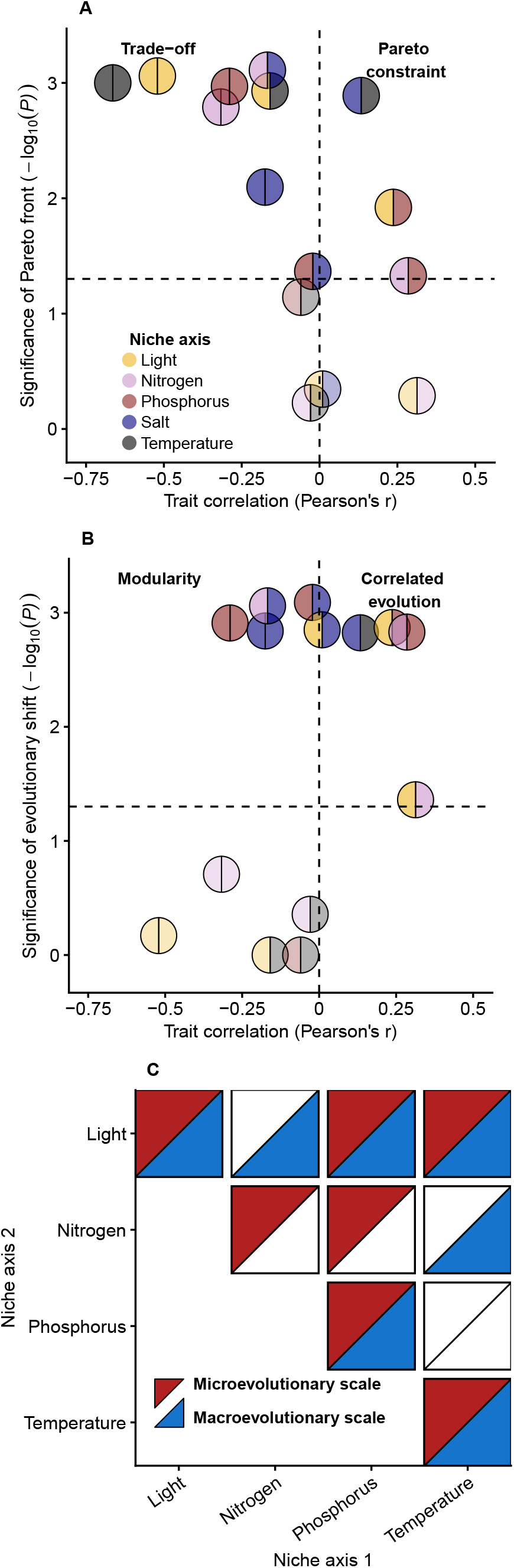
Summary of key results. (A) Relationship between the significance of Pareto fronts (*P*-values from one-tailed permutation tests) and trait correlations estimated with Pearson’s *r*. (B) Relationship between the significance of evolutionary shifts (P-values from linear mixed-effects models, one-tailed contrasts) within populations that evolved under matching environmental conditions (e.g. low-P tolerance in populations that evolved under limited phosphorus) and trait correlations. Points correspond to pairs of traits in a population (see Figures 2-3) and are coloured by niche axis (light, nitrogen, phosphorus, salt, temperature). Solid colour circles represent relationships within a single niche axis (*μ*_*max*_ vs. niche-determining trait), whereas split colour circles indicate relationships across distinct niche axes. Significant Pareto fronts or evolutionary shifts (*α* = 0.05, see horizontal dashed line) are shown in bolder colour; non-significant relationships are more translucent. (C) Comparison of the occurrence and significance of Pareto constraints across microevolutionary (experimentally evolved populations of *Chlamydomonas reinhardtii*) and macroevolutionary (189 phytoplankton species) scales. Coloured triangles indicate whether a significant Pareto front is detected for each trait pair in each dataset: filled triangles denote a significant front, white triangles indicate no statistical support for a Pareto front.

### Micro- and macroevolutionary trade-offs differ across phytoplankton

The patterns of phenotypic constraint we observed in experimentally evolved populations of *Chlamydomonas reinhardtii* did not always extend to macroevolutionary relationships among phytoplankton species. Among our *C. reinhardtii* populations, we observed consistently negative trait correlations between *µ*_*max*_ and niche-determining traits within individual niche axes (Figure 2). However, negative correlations were not evident across diverse phytoplankton taxa (Figure 4, Table S5). Despite the absence of these negative correlations at the macroevolutionary scale, we detected strong evidence of Pareto fronts constraining the joint optimization of *µ*_*max*_ and niche-determining traits within light, phosphorus, and temperature gradients across species (one-tailed permutation tests, *P* ≤ 0.022, Figure 4AHJ). These constraints closely mirrored those observed in experimental *C. reinhardtii* populations (*P* ≤ 0.001, Figures 2ACE, 5C), suggesting that while evolutionary diversification and innovation can relax and overcome trait correlations, underlying physiological limits on growth and resource use optimization persist across phytoplankton.

In contrast to comparisons within niche axes, relationships between niche-determining traits across niche axes were rarely consistent across evolutionary scales. For most relationships, we detected Pareto fronts only at one evolutionary scale (Figure 5C). Among experimentally evolved populations, we detected a significant Pareto front between low-N and low-P tolerance (*P* = 0.047, Figure 3E) that was absent in our cross-species analysis. Conversely, we detected significant Pareto fronts in the low-N-low-L tolerance and low-N tolerance-thermal breadth relationships across phytoplankton (*P* ≤ 0.005, Figure 4BG) but did not observe them within our experimentally evolved populations (Figure 5C). We found consistent evidence for Pareto constraints across micro- and macroevolutionary scales in only 2 of 6 relationships across niche axes (low-L-low-P tolerance and low-L tolerance-thermal breadth, *P* ≤ 0.012, Figure 5C). Overall, significant Pareto fronts were shared across micro- and macroevolutionary scales in 5 of 10 relationships (Figure 5C)—3 of 4 within single niche axes, but only 2 of 6 across niche axes. Notably, Pareto fronts emerged even in the absence of significant negative trait correlations at both scales—universally across species (Figure 4), and for several fronts among our experimentally evolved populations (Figure 3).

## Discussion

In this study, we demonstrate widespread Pareto fronts constraining relationships between growth rates and key niche-determining traits including thermal breadth, low-resource tolerance, and salt tolerance in phytoplankton. These constraints are evident at both microevolutionary scales (in experimentally evolved populations of *Chlamydomonas reinhardtii*) and macroevolutionary scales (across 189 species), highlighting the central role of trade-offs in shaping phenotypic evolution. Critically, such trade-offs do not necessarily manifest as simple overall negative correlations among traits (3, 5, 21) but instead emerge most clearly among Pareto-optimal genotypes. By defining the upper bounds of performance, Pareto fronts delineate regions of phenotypic space that are effectively inaccessible, consistent with the persistent emptiness of large portions of theoretically possible phenotypic space across the tree of life (35, 42, 43).

Within phytoplankton communities, trade-offs among traits associated with nutrient acquisition and utilization, growth, and cell morphology are widely implicated in driving ecological divergence and maintaining biodiversity (23). Here, we find strong evidence for trade-offs between niche-determining traits and maximum growth rates that recapitulate classic gleaner-opportunist trade-offs in nutrient acquisition (20-22), generalist-specialist trade-offs in thermal traits (26), and salt tolerance-growth trade-offs (24, 25). In our study, these trade-offs—manifesting as negative correlations between traits—consistently emerged among experimentally evolved *C. reinhardtii* populations (Figure 5A, Table S2). This pattern is consistent **with** ecological trade-offs in phytoplankton arising from evolutionary constraints such as antagonistic pleiotropy or resource allocation trade-offs (13, 23, 54), which can generate negative trait correlations not readily overcome through short-term adaptive evolution.

Despite their central role in ecology and evolution (4, 8, 11, 23, 27, 29, 45, 47), trade-offs can often be difficult to detect empirically, even for traits expected to incur substantial costs (31-33). When organisms differ widely in overall quality or resource budgets, high-performing individuals can excel across multiple tasks and traits at once, generating positive correlations that mask underlying trade-offs and reveal them only among the most optimized genotypes (3, 33). By explicitly characterizing the outer boundaries of achievable trait combinations rather than average relationships among them, Pareto fronts can recover constraints overlooked by correlational approaches and by more flexible measures of statistical dependence (e.g. mutual information), which capture linear and non-linear associations but likewise summarize entire trait distributions (61, 62). Pareto fronts thus allowed us to detect trade-offs in our experimentally evolved *Chlamydomonas reinhardtii* populations even where underlying trait correlations were positive or neutral (Figure 3). For example, we detected strong Pareto constraints on the joint optimization of low-phosphorus tolerance with both low-light and low-nitrogen tolerance despite significant positive correlations between these traits (Figure 5A). These results help reconcile observations of convergent evolution towards increased competitiveness for multiple essential nutrients (21, 30) with theoretical and empirical explorations of the importance of trade-offs in structuring communities (11, 23). Behind Pareto fronts that define intractable limits to multivariate trait optimization, adaptive evolution can generate positive correlations between traits, illustrating how evolutionary constraints on organismal performance can exist alongside adaptive trait correlations (3).

In our experimentally evolved *C. reinhardtii* populations, genotypes were generally only able to optimize multiple niche-determining traits simultaneously when the underlying correlations between traits were neutral or positive (Figure 5B); negative correlations were instead more often associated with limited movement towards Pareto optimality. These patterns suggest that the structure of phenotypic covariance behind a Pareto front strongly influences whether evolution can approach an optimal trade-off surface or remains constrained by genetic architecture prior to reaching it (13, 42, 43). Negative correlations did not, however, always prevent optimization. For the two most evolutionarily responsive traits—low-P and salt tolerance—selection under phosphorus limitation and salinity stress moved populations towards growth-tolerance Pareto fronts regardless of negative correlations (Figure 2CD), whereas less labile traits such as low-light and low-nitrogen tolerance showed little movement (Figure 2AB). Trait optimization in the face of negative correlations likely reflects modular or independent changes. While mutations with large, positively pleiotropic effects may be favoured when populations exist away from Pareto fronts, modular mutations of smaller effect are likely to become more common as phenotypes approach these limits **(43, 63-66)**.

Salt tolerance provided the clearest example of modular optimization: it increased consistently under salinity stress without detectable antagonistic effects on other niche-determining traits, allowing populations to approach Pareto fronts across multiple niche axes irrespective of negative or neutral trait correlations (Figure 5B). The unique modularity of evolved salt tolerance in our data likely reflects the novel intensity of this stressor relative to typical conditions experienced by *C. reinhardtii*—a species that lives in soil and freshwater—and is consistent with its capacity to evolve dramatically increased salt tolerance (67, 68). It also likely reflects the physiological distinctness of salt tolerance: tolerating high salt relies heavily on osmoregulatory processes involving the accumulation of compatible solutes and phospholipid metabolism (68, 69), largely separate from the core metabolic remodelling and upregulation of membrane transport proteins associated with limited nutrient acquisition (30), and the central role of heat-shock proteins in governing thermal tolerance (70, 71) in phytoplankton. Even so, each of these processes is energetically costly, drawing on shared energy budgets and overlapping machinery, including the production of membrane transport proteins. We therefore expect that trade-offs would ultimately manifest across all niche-determining traits in phytoplankton, even if they are weaker for comparatively modular, highly evolvable traits like salt tolerance (67, 68).

Strikingly, several patterns observed in our experimentally evolved *Chlamydomonas reinhardtii* populations did not translate directly to macroevolutionary patterns across phytoplankton species. In particular, the negative correlations between growth rate and niche-determining traits that were pervasive within individual niche axes at the microevolutionary scale (Figure 2) were absent across diverse phytoplankton taxa (Figure 4). Nonetheless, we found strong evidence that performance within these niche axes remains bounded by Pareto fronts at both evolutionary scales. Across 189 phytoplankton species, Pareto fronts significantly constrained the joint optimization of growth rate and niche-determining traits under light and phosphorus limitation and across thermal gradients, closely mirroring the boundaries observed in experimentally evolved *C. reinhardtii* populations (Figure 5C). The persistence of these limits on performance, despite the relaxation of genetic constraints over macroevolutionary timescales (54–57) and ample opportunity for evolutionary innovation, suggests that some trade-offs reflect fundamental physiological or ecological limitations rather than transient genetic constraints.

Relationships among niche-determining traits across niche axes were even less consistent between micro- and macroevolutionary scales. Several trait combinations that were constrained within experimentally evolved *Chlamydomonas reinhardtii* populations did not exhibit comparable Pareto front patterns across phytoplankton species, while others appeared constrained only at the macroevolutionary scale (Figure 5C). These discrepancies likely reflect the fundamentally different evolutionary processes operating across timescales: while experimental evolution captures short-term responses within a shared genetic and physiological background, macroevolutionary patterns emerge through long-term diversification across lineages occupying distinct ecological niches (5). Over deep evolutionary time, major innovations in phytoplankton biology—including coloniality and mixotrophy (50–52)—may alter or relax the genetic and physiological constraints that shape trait relationships within populations. Notably, these macroevolutionary Pareto fronts emerged despite the complete absence of significant negative trait correlations across phytoplankton (Figure 4). As at the microevolutionary scale (Figures 2-3), Pareto fronts revealed constraints on trait optimization that correlational approaches would have missed. Broadly, these results suggest that Pareto fronts can be a powerful tool for detecting trade-offs across biological scales—and help reconcile the long-standing difficulty of documenting trade-offs empirically with their fundamental importance in evolutionary ecology (2, 3, 5, 31, 33).

Some of the constraints we observed in our data likely reflect trade-offs involving more than two traits. Across phytoplankton, phenotypic distributions were often roughly triangular (Figure 4), consistent with multivariate selection; when selection acts on three or more traits, accessible phenotypes form a polygonal constraint envelope whose vertices are specialized archetypes and whose edges are the Pareto fronts that optimally balance pairs of traits (10, 24, 35). Each bivariate front we identified can thus be considered a single edge of such a polygon—the optimal trade-off surface for two traits, nested within a higher dimensional constraint envelope. In phytoplankton, gains in low-resource tolerance may incur costs in other key traits such as cell size (27), defence (28), or thermal flexibility (44, 72), helping to maintain apparently sub-optimal variation in niche-determining traits (Figure 1D). In our experimentally evolved *C. reinhardtii* populations, our focal traits traded off directly against cell size and pigmentation (Figures S6, S7), providing evidence for the sort of multivariate trade-off that may help explain the approximately triangular distributions we sometimes observed (Figure 3). That experimental populations nonetheless often converged on individual bivariate fronts may reflect the focused selection we imposed—stressing one or two niche axes at a time may drive populations toward the corresponding edge of a polygon, whereas the more multivariate fitness landscape of natural habitats may instead favour interior, generalist phenotypes that balance a full suite of traits (35, 42).

The trade-offs we observed reflect the imprint of past adaptation (12, 42, 43), but they will also shape—and constrain—how phytoplankton respond to the multivariate stressors of global change, including warming, salinization, and shifting nutrient regimes (6, 73). When several stressors act at once, Pareto fronts will limit how far populations can improve performance across all relevant dimensions (14–16), and facilitative correlations will only aid adaptation until populations reach these boundaries (Figure 5B). Our results indicate that such limits are not merely theoretical: we detected widespread Pareto fronts at both the population (Figures 2-3) and species (Figure 4) levels in phytoplankton. Both adaptive evolution within species and community turnover across species are thus expected to unfold within persistent performance boundaries. Understanding and predicting biological responses to global change therefore requires explicitly accounting for these limits on multivariate trait optimization rather than treating responses to individual stressors in isolation.

In this study, we demonstrate that widespread Pareto fronts—trade-offs that emerge among optimized phenotypes—constrain the evolution of niche-determining traits across both micro- and macroevolutionary scales in phytoplankton. By extending Pareto fronts as an analytical framework for detecting context-dependent and otherwise elusive trade-offs (3, 5, 31), our results highlight their utility for understanding phenotypic evolution under constraint at multiple evolutionary timescales. Across experimental evolution and cross-species comparisons, we show that adaptation is not solely limited by genetic variation or opportunity, but also by fundamental, multivariate limits on trait optimization. Recognizing and characterizing these limits is therefore essential for understanding how phenotypic diversity is generated, maintained, and constrained in a changing world.

## Materials and Methods

### Experimental evolution of *Chlamydomonas reinhardtii*

We used experimentally evolved populations of *Chlamydomonas reinhardtii* (21, 30) to examine phenotypic trade-offs among niche-determining traits. In brief, we inoculated experimental chemostats with *C. reinhardtii* cells cultured from one of four ancestral isoclonal colonies and from a genetically diverse original population. We evolved each ancestral population under conditions of salt stress (N = 5), light (N = 5), phosphorus (N = 5), and nitrogen (N = 5) limitation, and in biotically depleted media (N = 5), a biotic depletion × salt stress interactive treatment (N = 4), and a control COMBO (74) treatment (N = 3). We experimentally evolved *C. reinhardtii* populations for 285 days. Though we started with 5 replicate populations in each evolution treatment (N = 5 per treatment), 3 of the 35 evolved populations were lost to contamination, resulting in 32 descendant populations which we isolated for subsequent common garden experiments. We then used four independent replicates from each population in common garden batch-culture experiments to assess their niche-determining traits (low-resource tolerance for light (1/*L* *), nitrogen (1/*N*\*), phosphorus (1/*P*\*), and thermal breadth (*T*_*br*_), though no populations were evolved under thermal stress), and a single replicate for salt tolerance (S). Full details of our experimental design are available in the Supporting Methods (2.i-ii).

### Growth assays and estimation of niche-determining traits

We quantified *Chlamydomonas reinhardtii* growth across gradients of light, nitrogen, phosphorus, salinity, and temperature, generating high-resolution growth time series. We fit growth rate data with thermal performance curves, Monad functions curves, Monod functions, and inverse logistic growth models to estimate niche-determining traits (Figure S1), including maximum growth rate (*μ*_*max*_), low-resource tolerance for light, nitrogen, and phosphorus (1/*R*^*\**^), salt tolerance (*S*), and thermal breadth (*T*_*br*_). Full details of experimental design, model selection, and fitting procedures are provided in the Supporting Methods (2.iii-iv).

### Detection and statistical evaluation of Pareto fronts

To identify trade-offs constraining multivariate trait optimization, we analyzed bivariate relationships among niche-determining traits to detect putative Pareto fronts—upper boundaries of phenotypic distributions beyond which simultaneous optimization of multiple traits is not observed (42, 43). For each bivariate relationship, we identified the set of Pareto-optimal phenotypes—the points on the upper right convex hull for which no other observation achieved higher values for both traits—and fit a smoothed, monotonically decreasing curve to these points to visualize the front (with curvature unconstrained). To quantify patterns of constraint, we used the convex hull to calculate the area of empty, inaccessible phenotypic space in the upper right corner of each plot (shaded area in Figure 1C). We assessed the significance of each putative Pareto front using a one-tailed permutation test (42), wherein we compared the observed inaccessible area to a null distribution generated by independently randomizing the values of each trait 1,000 times. We calculated *P* as the proportion of randomizations in which the inaccessible area equaled or exceeded the observed value. Because a partial convex hull—and thus a putative Pareto front—can be fit to nearly any bivariate distribution, including randomly scattered data, we considered only cases where the observed inaccessible space exceeded null expectations (*α* = 0.05) as providing evidence for the existence of Pareto fronts. Non-significant fronts (shown in grey in Figures 3-4) are therefore displayed for reference only and provide no evidence of evolutionary constraint. We note that the traits in our models were all estimated with error that was not propagated in our Pareto front analyses. Because random estimation error tends to blur, rather than create, the boundaries of trait distributions, this likely renders our tests of Pareto front significance conservative. Additional details are provided in the Supporting Methods (2.vi-viii).

The location of a Pareto front is set by the boundary of a trait distribution and is unchanged by interior points—the front is identical regardless of how many sub-optimal phenotypes lie behind it. These sub-optimal points do, however, affect the probability of detecting a significant Pareto front, because they contribute large numbers of low trait values to the null randomizations and inflate the size of the null inaccessible areas. We therefore assessed Pareto fronts using the third of phenotypes furthest from the minimal (x, y) values (and closest to the Pareto front), excluding points that were uninformative about the front’s location while reducing our power to detect it. A sensitivity analysis (Supporting Methods 2.ix) showed that our results were robust to other threshold levels: 8 of the 11 Pareto fronts significant at a 33% data retention threshold (including all within-axis fronts) remained significant across thresholds from 25% to 100% (Figure S2, Table S1). The remaining three, weaker fronts (all involving low-P tolerance) were significant only at more stringent thresholds (≤ 33% retention). The detection of Pareto fronts only became unreliable at the extremes—when too few observations remained (10%), or when large numbers of sub-optimal interior points obscured weaker Pareto fronts.

To compare our Pareto fronts against a traditional, correlational assessment of trade-offs, we estimated the Pearson correlation (*r*) for every pair of traits. When correlations were significant, we visualized these relationships using standardized major axis (SMA) regression lines, an appropriate method for summarizing bivariate relationships when neither variable is a natural predictor of the other (75).

To test whether experimental evolution moved populations toward Pareto-optimal trait combinations, we asked whether populations that evolved under conditions matching a given niche axis came to lie closer to the corresponding Pareto front than their ancestors. For each bivariate relationship, we z-scored both traits, constructed a stepwise (un-smoothed) Pareto front through the Pareto-optimal phenotypes, and calculated the Euclidean distance in this z-scored trait space from each replicate population to its nearest point on the front (blue and grey dashed lines in Figures 1C, S3). We used linear mixed-effects models (distance ∼ evolutionary history + (1(ancestry)) with one-tailed contrasts to test each matching treatment separately, asking whether populations that evolved under the matching environment (e.g. salt-stressed populations for the *µ*_*max*_-S relationship) lay significantly closer to the Pareto front than ancestral populations (i.e. comparing the lengths of blue (descendant) vs. grey (ancestral) dashed lines in Figure 1C). Full details for the front construction, distance calculation, and model specification are provided in the Supporting Methods (2.xi).

### Synthesis and analysis of phytoplankton trait variation across species

To assess whether constraints observed in our single species evolution experiment (*C. reinhardtii*) extend across broader macroevolutionary scales, we compiled published estimates of phytoplankton growth rates measured across gradients of light, nitrogen, phosphorus, and temperature (60, 72, 76-80). From these data, we fit thermal performance and Monod curves to estimate niche-determining traits and screened all fits for quality, yielding 499 resource- or temperature-dependent growth models across 189 phytoplankton species. Because some species contributed multiple data points, we averaged trait estimates within each species prior to analysis. We then analyzed bivariate trait relationships across species using the convex hull-based Pareto front methods, empty-space randomization tests, and correlational approaches described above. Full details on data compilation, model fitting, and statistical analyses are provided in the Supporting Methods (2.xiii-xiv).

## Data, materials, and software availability

All code and data required to replicate analyses and generate figures are archived at Zenodo (DOI: 10.5281/zenodo.19032401) and available via GitHub (https://github.com/JasonLaurich/Chlamy_37_pops_pheno_constraint).

## Acknowledgments

We thank Daniel Steiner for assistance with HPLC analysis, Ken Thompson for manuscript feedback, Megan Szojka for code review, as well as Van Savage and two anonymous reviewers for their constructive reviews and feedback. Our work was supported by a Natural Sciences and Engineering Research Council of Canada (NSERC) Discovery Grant and a Canada First Research Excellence Fund Grant awarded to J.R.B. J.R.B.’s work during the lab experiments phase of this study was supported by a Nippon Foundation Nereus Postdoctoral Fellowship. We also thank members of the Bernhardt Lab for helpful feedback on an earlier version of this manuscript.

## Notes

https://zenodo.org/records/21987486

### Competing Interest Statement

The authors have declared no competing interest.

### Summary of Updates

This version has been heavily revised following 2 rounds of peer review. Major changes include improvements to figure clarity, streamlining of analyses to reduce complexity, and textual elaboration on key themes and concepts.

