## Supporting text for "Pareto fronts reveal constraints on the evolution of ecological niches"

### Supporting information for
Pareto fronts reveal constraints on the evolution of ecological niches

Jason R. Laurich[1], Anita Narwani[2], & Joey R. Bernhardt[1]

[1]Department of Integrative Biology, University of Guelph, Guelph, ON, Canada

[2]Department of Aquatic Ecology, Swiss Federal Institute of Aquatic Science and Technology (Eawag), Dübendorf, Switzerland

*Corresponding author: Jason R. Laurich

**Author Contributions:**

Conceptualization: J.R.L., A.N., J.R.B. Data curation: J.R.L., A.N., J.R.B. Formal analysis: J.R.L. Funding acquisition: A.N., J.R.B. Investigation: J.R.L., J.R.B. Methodology: J.R.L., A.N., J.R.B. Project administration: J.R.L., A.N., J.R.B. Resources: A.N., J.R.B. Software: J.R.L. Supervision: J.R.B. Validation: J.R.L., J.R.B. Visualization: J.R.L. Writing—original draft: J.R.L. Writing—review & editing: J.R.L., A.N., J.R.B.

**Competing Interest Statement:** The authors have no competing interests to declare.

**Classification:** Biological Sciences/Evolution

**Keywords:** Pareto fronts, evolutionary constraint, trade-offs, ecological limits, phytoplankton

**This PDF file includes:**

Table of Contents
Part A: Supporting Text
Part B: Supporting Figures 1 to 8
Part C: Supporting Tables 1 to 10
References

### Part A: Supporting Text

#### 1. Supporting Results

**i. Principal component analysis and redundancy analyses identify trade-offs in multivariate phenotypic space.**

Principal component analysis of niche-determining traits, alongside pigmentation, stoichiometry, and cell size, revealed consistent trade-offs between maximum growth rates ($\mu_{max}$) and corresponding niche-determining traits within individual niche axes. Within light, nitrogen, phosphorus, salt, and temperature gradients, growth rates and their associated niche traits loaded in opposing directions along one or more principal component axes (Figure S6), consistent with negative relationships observed in Pearson correlations and Pareto front analyses (Figure 2).

In our analysis, the first two principal components explained 22.77% and 17.36% of variation, respectively. In addition to the trade-offs between growth and niche-determining traits above, we observed strong positive covariance among niche-determining traits that recapitulated positive correlations (e.g. the $1/L^*$-$1/P^*$ and $1/N^*$-$1/P^*$ relationships, Figure 3BE). Variation in pigmentation (chlorophyll *a*, chlorophyll *b*, and lutein) was also highly collinear.

To disentangle the relative contributions of ancestry and evolutionary environment to overall variation in growth rates, niche-determining traits, pigmentation, cell size, and stoichiometry in our experimentally evolved *C. reinhardtii* populations, we performed redundancy analyses (RDA). We found that experimental evolutionary environment consistently explained more total multivariate phenotypic variation than ancestry (23.18% vs. 12.53%, Figure S5AC). In both cases, explained variation was dominated by contributions from cell size, and cellular nitrogen and phosphorus content. After removing these variables, the relative importance of evolutionary environment became even more pronounced, explaining 25.86% of total variation compared to only 2.06% explained by ancestry (Figure S5BD).

**ii. Sensitivity analysis and selection of data retention threshold.**

The location and shape of a Pareto front is determined by the upper boundary of trait variation rather than the distribution of internal points. Because these interior or sub-optimal data points do, however, affect the statistical probability of detecting large empty spaces in null, randomized data sets, we performed a sensitivity analysis to interrogate the consequences of thresholding our data to include only relatively optimized data at six thresholds (10%, 25%, 33%, 50%, 67%, and 100% of the data).

Critically, our key findings—that Pareto fronts act as constraints on multivariate trait optimization and generate trade-offs not visible through traditional correlational approaches—were qualitatively consistent across a broad range of thresholds (25-100% data retained), breaking down only at the most stringent threshold (10%), where few data points remained (Figure S2, Table S1). Of the 11 Pareto fronts significant at the 33% threshold, 8 (including all of our single-gradient Pareto fronts) remained significant across all intermediate thresholds (25-100% data retained); only at the most extreme 10% retention did the $\mu_{max}$-salt tolerance and low-N tolerance-salt tolerance fronts lose significance, reflecting the reduced statistical power of so few remaining data points (Figure S2, Table S1). Retaining only 10% of the data also produced significant fronts in two relationships (low-L-low-N tolerance and low-P tolerance-thermal breadth, Figure S2BL) that we did not detect at any other threshold, suggesting that so few observations make the determination of statistical significance unstable in both directions. At the other extreme, retaining more sub-optimal points (e.g. relaxing the data retention threshold from 33% to 50% retained) inflated null expectations and obscured the three weaker fronts (low-P-low-L tolerance, low-P-low-N tolerance, and low-P-salt tolerance, Figure S2CGK), reducing the number of significant Pareto fronts in our analysis from 11 to 8 (of 15 relationships). We therefore treat intermediate thresholds (25%-67%) as the most reliable range for estimating the significance of Pareto fronts, and interpret these three weaker Pareto fronts with caution (see main text, Results).

**iii. Pareto fronts between niche-determining traits, pigmentation, biovolume, and stoichiometry in *Chlamydomonas reinhardtii*.**

We examined multivariate relationships between niche-determining traits (1/*L**, 1/*N**, 1/*P**, salt tolerance, and thermal breadth) and additional physiological traits, including cell size (biovolume), elemental stoichiometry (cellular N and P content), and pigmentation. Because chlorophyll *a*, chlorophyll *b*, and lutein were highly collinear (Figure S6), we summarized pigmentation using scores along the first principal component of a pigment-only PCA.

Pareto front analyses revealed many constraints between niche-determining traits and both biovolume and pigmentation (Figure S7), with inaccessible phenotypic space significantly exceeding null expectations for all niche-determining traits except the 1/*P**-pigmentation, 1/*P**-biovolume, and $T_{br}$-pigmentation relationships. We also detected additional Pareto fronts in the $T_{br}$-P content and salt tolerance-P content relationships (Figure S7NR, Table S6).

### 2. Supporting Materials and Methods

All analyses and figures in this study can be reproduced by running the master script 00_master_script.R, which sources the individual scripts that perform particular analyses and are referenced throughout the Supporting Materials and Methods. The computationally intensive scripts (Bayesian model fitting, scripts 01-05 and phytoplankton synthesis data model fitting, scripts 14-17) are commented out by default so that the master script runs quickly from our processed data. All code and data are archived at Zenodo (DOI: 10.5281/zenodo.19032401).

**i. Experimental evolution.**

In February 2015, we isolated four ancestral colonies grown from single *Chlamydomonas reinhardtii* (cc1690 wild-type mt+) cells obtained from the Chlamydomonas Resource Centre (chlamycollection.org) from agar plates. We experimentally evolved these ancestors, as well as mixed-ancestry populations collected from the original cc1690 population over 285 days (~285 generations) in continuous flow chemostats. We maintained chemostats with 28 mL of experimental media, and replenished media at a rate of 56% per day with a peristaltic pump. Chemostats were also continuously mixed and aerated to ensure equal distribution of dissolved resources. We supplied all chemostats with modified COMBO media (1) lacking silicon, vitamins and animal trace elements and maintained them at 20°C under 90 µmol photons $m^{-2}s^{-1}$ of light (except for the light limitation treatment) on an 18:6-hour light:dark cycle. The control (C) treatment received full strength COMBO solution (1) supplied with 1000 µM N and 50 µM P. To experimentally evolve *Chlamydomonas reinhardtii* under conditions of nutrient limitation, we reduced concentrations of relevant nutrients (nitrogen, N: 10 µM, phosphorus, P: 0.5 µM). In our light limitation treatment (L) we reduced illumination to 5 µmol photons $m^{-2}s^{-1}$ using neutral-density light filter paper (Solar Graphics™, Clearwater, Florida), and supplemented control COMBO with 8 g/L NaCl for our salt treatment (S). We prepared our biotically-depleted medium (B) by culturing seven other freshwater algae in control COMBO (*Cosmarium botrytis*, *Kirchneriella subcapitata*, *Pediastrum boryanum*, *Pediastrum duplex*, *Scenedesmus acuminatus*, *Staurastrum punctulatum*, and *Tetraedron minimum*) individually. We removed algal biomass from these cultures via filtration and autoclaved media to ensure sterility. These seven sterilized filtrates were then mixed to create our biotically-depleted media (B) and stored at 4°C in the dark prior to use in experiments. We added 8 g/L NaCl to this medium to create the combination salt stress and biotic depletion treatment (BS). Across all experimental treatments, we ramped up the severity of nutrient limitation, salt stress, light limitation, and the effects of biotic depletion each month, starting with full-strength control COMBO before gradually reaching final experimental conditions (Table S7). After 285 days, we recovered descendant populations and inoculated them onto fresh agar plates for long-term storage. We evolved each of the five ancestral populations under all seven treatments (35 experimental chemostats). In total, 32 descendant populations were collected from the 5 ancestral populations while three populations (C–ancestors 2 and 3, BS-ancestor 5) were lost to contamination over the course of experimental evolution. Together with the 5 ancestral populations, this left us with 37 experimental populations of *C. reinhardtii* for the common garden experiments.

**ii. Assessing minimum resource requirements, salt tolerance, and thermal performance.**
We conducted batch culture experiments with all 32 descendant populations of *Chlamydomonas reinhardtii* and the five ancestral populations (mixed population and isoclonal Ancestors 2-5) to quantify niche-determining traits. We estimated minimum nitrogen, phosphorus, and light requirements ($N^*$, $P^*$, and $L^*$, defined as the minimum resource level required for positive population growth), salt tolerance (*S*, defined as the salinity at which populations reached half their maximum growth rate), and thermal performance. To estimate $N^*$, $P^*$, $L^*$, and *S*, we measured population growth rates across ten levels of each abiotic gradient (Table S8). We conducted growth rate experiments in the inner 60 wells of 96-well plates, with each well containing 125 µL of modified COMBO medium inoculated with a low-density population (~10 relative fluorescence units, RFUs). We replicated each population × treatment four times, except for salt experiments, which were not replicated.

To minimize carry-over effects from stock culture conditions, populations were acclimated prior to growth assays. For nitrogen, phosphorus, and light limitation experiments, we acclimated populations at both the lowest resource level and at half the maximum resource level. We used cultures acclimated at moderate resource levels to inoculate high-resource treatments, and used cultures acclimated at low resource levels for treatments at or below half the maximum resource concentration. For salt tolerance assays, we acclimated populations at five salinity levels (0, 2, 4, 6, and 8 g $L^{-1}$ NaCl) and inoculated experimental treatments using cultures acclimated to the nearest lower salinity.

We filled the outer wells of each plate with COMBO medium to reduce evaporative loss, and plates were sealed with gas-permeable Breathe-Easy membranes (Sigma-Aldrich). We quantified population growth using chlorophyll *a* fluorescence, measured as relative fluorescence units (RFUs), using a Biotek Cytation 5 plate reader (excitation: 435 nm; emission: 685 nm). For nitrogen, phosphorus, and light limitation assays, we measured RFUs two to three times daily over three days to capture exponential growth. For salt tolerance assays, we measured RFUs once daily for nine days. We assessed thermal performance by measuring growth at six temperatures (10, 16, 22, 28, 34, and 40°C; Table S8), with four replicates per population at each temperature. We measured RFUs two to three times daily, depending on the relative growth rates of *C. reinhardtii* at each temperature. We did not acclimate populations at experimental temperatures prior to the experiments, allowing us to quantify acute thermal responses across the temperature gradient.

**iii. Estimating population growth rates.**
We carried out our mathematical model fitting and subsequent analyses in R version 4.4.1 (2)**.** All code and data required to replicate all analyses and generate all figures for this study are archived at Zenodo (DOI: 10.5281/zenodo.19032401). We estimated the population growth rates of *Chlamydomonas reinhardtii* replicates from changes in relative fluorescence units (RFUs) over time during the exponential growth phase at each level of each abiotic gradient. For each replicate time series, we identified the exponential growth phase using a sliding-window approach that fit log-linear models to each successive subset of log-transformed RFU data spanning the full time series. We took the window yielding the maximum log-linear slope to represent the exponential growth period. We modeled growth rate during the exponential phase as:

$$F(t) = F(0)e^{\mu t} \quad (1)$$

using nonlinear least squares implemented in the nls.multistart package in R (3), where $F(t)$ is the measure of RFUs at time *t*, $F(0)$ is the replicate-level RFU measurement at time 0, and $\mu$ is the exponential growth rate of the population in $days^{-1}$ (4).

The code responsible for estimating exponential growth rates can be found in the following scripts: 01_light.R, 02_nitrogen.R, 03_phosphorus.R, 04_salt.R, and 05_temperature.R.

**iv. Estimating niche-determining traits.**

We estimated thermal performance curves (TPCs; Figure S1A-B) for each replicate using exponential growth rates measured across temperature gradients. We modeled thermal performance curves using Bayesian hierarchical models implemented in R2jags (5), interfacing with JAGS. For each population, we fit models predicting growth rate ($day^{-1}$) as a function of temperature. All models were run with six independent Markov chain Monte Carlo (MCMC) chains, each iterated for 330,000 steps with a burn-in of 30,000 and thinning interval of 300, yielding 6,000 posterior samples per model. We assessed model convergence by using trace plots and ensuring that $\hat{R}$ values were approximately 1 for each model parameter. We also confirmed model independence by ensuring that effective sample sizes exceeded 60% of posterior samples for all parameters.

To identify an appropriate functional form for TPCs, we compared 12 candidate models commonly used to describe ectotherm thermal performance using Akaike Information Criterion (AIC) as implemented in the rTPC package (6, 7). Based on model performance across populations, we selected the Lactin II model (8), which consistently provided good fits and allows growth rates to become negative at extreme temperatures (Table S9). The Lactin II model describes growth rate as a function of temperature:

$$\mu(T) = e^{\rho T} - e^{\rho T_{max} - \left(\frac{T_{max} - T}{\Delta T}\right)} + \lambda \qquad (2)$$

where $\mu(T)$ is a biological rate (e.g. growth rate) at temperature $T$, $\rho$ is a temperature sensitivity parameter that models the rate of increase at sub-optimal temperatures, $T_{max}$ is the optimal temperature beyond which performance sharply declines, $\Delta T$ controls the steepness of this decline, and $\lambda$ shifts the intercept.

We fit Lactin II models to growth–temperature data for each replicate and derived biologically meaningful thermal traits from the fitted curves. Using the posterior distributions of our models, we calculated thermal optima ($T_{opt}$) by numerically solving for the temperature at which the first derivative of the fitted curve equals zero using the Deriv package (9) and uniroot functions in R. We calculated maximum growth rate ($\mu_{max}$) by evaluating the fitted model at $T_{opt}$. We identified minimum ($T_{min}$) and maximum ($T_{max}$) temperatures supporting positive growth as the lower and upper roots where $\mu(T) = 0$. We calculated thermal breadth ($T_{br}$) as the temperature range over which growth exceeded 50% of $\mu_{max}$, calculated as the difference between the upper and lower roots of $\mu(T) = \mu_{max}/2$. We calculated each estimate for each independent Bayesian posterior and report the median values for each. We used moderately informative priors, shared across populations, for all TPC parameters (Table S10).

We fit Monod curves (Figure S1D-F) to exponential growth rates across abiotic gradients for nitrogen, phosphorus, and light limitation by modeling resource-dependent growth rates, $\mu(R)$ as:

$$\mu(R) = \mu max\left(\frac{R}{k_s + R}\right) \qquad (3)$$

where $\mu_{max}$ is the maximum exponential growth rate, $R$ is the resource level, and $k_s$ is the half-saturation constant, the resource concentration at which growth is one half of $\mu_{max}$ (10). For all replicates and populations, we set moderately informative priors that worked for all three nutrient gradients (Table S10) and confirmed model performance as described above.

We calculated minimum resource requirements ($R^*$) for nitrogen, phosphorus, and light limitation ($N^*$, $P^*$, and $L^*$ respectively) using our parameter estimates from the Monod curve following:

$$R^* = \frac{m\, k_s}{\mu max - m} \tag{4}$$

where $m$ is the mortality rate, which we set to 0.56 day$^{-1}$ to match the mortality rate imposed in our chemostat experiments (4).

We defined salt tolerance as the salt concentration at which population growth rates were equal to half their maximum growth rate (4), and modelled the relationship between growth rate and salt concentration using a reverse logistic function (Figure S1G-I):

$$\mu([\mathrm{S}]) = \frac{a}{1 + e^{b([\mathrm{S}]-S)}} \tag{5}$$

where $a$ is the maximum population growth rate, $b$ models the decline in growth rate as salt concentration ([S]) increases, and $S$ is the salt concentration at which growth rates are half their maximum, or a population's salt tolerance. For our salt tolerance curves, we set broader priors to account for the strong effects of evolution under salt stress on salt tolerance and confirmed model performance as described above.

To explore and confirm the validity of our parameter estimates and model fits, we created a Shiny app that plots raw growth data and fitted model estimates for each of our experimental *C. reinhardtii* populations and replicates across temperature, light, nitrogen, phosphorus, and salinity gradients. The code required to recreate this app is archived in the 'Shiny-apps' folder at Zenodo (DOI: 10.5281/zenodo.19032401).

The code responsible for fitting Bayesian models to variation in exponential growth rates across abiotic gradients can be found in the following scripts: 01_light.R, 02_nitrogen.R, 03_phosphorus.R, 04_salt.R, and 05_temperature.R.

**v. Estimation of nitrogen and phosphorus content, cell size, and pigmentation.**

To quantify variation in cellular stoichiometry, we inoculated a single replicate of each population (ancestral or experimentally evolved) into 400 mL beakers with sterile COMBO medium (1). After approximately 1.5 days of growth (to reach the mid-exponential growth phase) we filtered each culture on one ashed (400°C) Whatman ® glass microfiber filter (grade GF/F, 47 mm) and one 25 mm Whatman ® glass microfiber filter. After drying the filters in a 60°C oven overnight, we weighed and calculated dry biomass of the samples by subtracting the mass of pre-weighed filters. We calculated the nitrogen content of algal biomass using the 47 mm filter in an Elementar vario PYRO cube EA-IRMS, and measured phosphorus content using the 25 mm filter in a Skalar San++ Continuous Flow P/N analyser. For our phosphorus samples, we first digested and oxidized samples using a peroxydisulphate solution, and diluted samples in a 1:20 ratio before running them on the P/N analyser.

After finishing the final RFU measurements in the batch culture experiments designed to quantify algal performance across light, nitrogen, and phosphorus gradients, we fixed cells by adding a 10% glutaraldehyde solution. We stored plates at 4°C before imaging them using a BioTek Cytation 5 imaging plate reader using the Brightfield setting. We analyzed cell length using Gen5 software (BioTek v. 2.0), which we converted to biovolume under the assumption of spherical cell shape. For analyses, we averaged observed biovolume measurements across all treatments, using only a single estimate of biovolume for each population. To measure the concentration of photosynthetic pigments (chlorophyll *a*, chlorophyll *b*, and lutein) we performed high-performance liquid chromatography (HPLC) using an HPLC system from Jasco Instruments (Easton, MD, USA).

**vi. Defining Pareto fronts.**

We quantified Pareto fronts in bivariate trait distributions to assess whether trade-offs among relatively optimized genotypes constrain realized phenotypic variation. Specifically, we examined relationships between maximum exponential growth rates ($\mu_{max}$) and niche-determining traits (minimum resource requirements, thermal breadth, and salt tolerance), as well as relationships among niche-determining traits themselves. Here we define Pareto fronts as the set of trait combinations representing optimal solutions to a trade-off, beyond which simultaneous improvement of both traits is not possible or exceedingly rare (11–13).

**vii. Identification of Pareto fronts.**

For each bivariate trait relationship, we identified candidate Pareto front points using a modified convex hull algorithm that retained only observations defining the upper-right boundary of trait space (12). These points represent phenotypes with jointly high values for both traits under consideration. We then fit monotonically decreasing, smoothed curves through these Pareto front points using shape-constrained additive models implemented in the scam package (14). To avoid overfitting, the maximum basis dimension was set to the number of Pareto front points, up to a maximum of six.

We applied this approach to test for classic ecological trade-offs, including gleaner–opportunist trade-offs between growth rate and competitive ability (4, 15), generalist–specialist trade-offs in thermal traits (16), and trade-offs between (salt) stress tolerance and growth (17, 18). We also examined relationships among niche-determining traits to evaluate whether adaptation to multivariate stressors is constrained by trade-offs.

The code responsible for assembling all of the niche-determining trait data into a single masterfile can be found in 06_data_compilation.R. The code for single gradient analyses can be found in 08_intra-gradient_trade-offs.R; cross-gradient analyses are found in scripts 09_inter-gradient_trade-offs.R and 10_secondary_trait_trade-offs.R.

**viii. Empty-space tests using randomized null models.**

To test the statistical significance of inferred Pareto fronts, we used an approach adapted from Li et al. (12) that quantifies the amount of phenotypic space rendered inaccessible by a front. For each fitted Pareto front, we calculated the area of empty space bounded by the front and the maximum observed values of the two traits using the polyarea function in the pracma package (Figure 1C; 19). We then generated 1,000 null datasets by independently randomizing x and y trait values, refit Pareto fronts to each null dataset, and recalculated the corresponding empty-space areas. We calculated *P* values with a one-tailed randomization test as the proportion of null datasets that produced empty-space areas equal to or larger than those observed in the real data.

Because only a subset of populations evolved under selective regimes relevant to any given trait combination, many observations in our dataset represent poorly optimized phenotypes located far below the Pareto front. These low-value points inflated null expectations and reduced statistical power in empty-space tests. To address this distortion, we performed all analyses on trimmed datasets from which the lower two thirds of observations were removed. We trimmed data points by z-scoring trait values and calculating the Euclidean distance of each observation relative to the minimum x and y values and removed the two thirds of data with the minimal scaled Euclidean distances.

The code responsible for fitting Pareto fronts and testing their significance for the experimentally evolved *C. reinhardtii* populations can be found in 11_Pareto_sensitivity_analysis.R.

**ix. Sensitivity analysis of Pareto front detection thresholds.**
Because the empty-space randomization test compares the observed inaccessible space to that expected under a null distribution, its power depends on the proportion of sub-optimal phenotypes included. Interior points that lie far below a front contribute low values to both x and y variables that inflate null expectations, making genuine constraints harder to detect. We therefore restricted our primary analyses to the third of phenotypes closest to each front, based on z-scored Euclidean distance from the minimum (x, y) value.

To confirm that this choice did not drive our conclusions, we repeated the full empty-space randomization tests across six data-retention thresholds—retaining 100%, 67%, 50%, 33%, 25%, or 10% of observations with the greatest scaled Euclidean distance from the minimum (x, y) values. At each threshold, we regenerated 1,000 null datasets based on the remaining data, recomputed a null distribution of inaccessible space area, and compared these estimates against our observed inaccessible space area to assess the significance of Pareto fronts (Table S1, Figure S2).

The code responsible for fitting Pareto fronts across data retention thresholds can be found in 11_Pareto_sensitivity_analysis.R.

**x. The correlational approach to measuring trade-offs.**
To contrast our Pareto front framework and results with a traditional, correlational view of trade-offs, we estimated the Pearson correlation (*r*) between each pair of traits and tested its significance using the cor.test function in R. For relationships with a significant correlation ($\alpha$ = 0.05), we fit standardized major axis (SMA) regressions using the smatr package (20) to visualize the bivariate relationship. Because SMA accounts for variation and error in the estimation of both traits rather than treating one as a predictor, it provides a symmetric summary appropriate for visualizing correlations between two traits measured with comparable uncertainty.

The code responsible for single gradient analyses can be found in 08_intra-gradient_trade-offs.R; cross-gradient analyses are found in scripts 09_inter-gradient_trade-offs.R and 10_secondary_trait_trade-offs.R.

**xi. Testing evolutionary shifts toward Pareto fronts.**
To test whether experimental evolution caused populations to move toward Pareto fronts in trait space, we implemented complementary bivariate and univariate approaches. For each bivariate relationship, we assessed whether populations evolved in a particular abiotic gradient (e.g. salinity stress) responded by optimizing the matching set of traits governing their performance across that same gradient (e.g. the $\mu_{max}$-salt tolerance relationship) relative to ancestral populations. For each relationship, we z-scored both traits, constructed a stepwise (un-smoothed) Pareto front using a convex hull algorithm, and calculated the Euclidean distance of each replicate from its nearest point on this front (Figures 1C, S3). We used linear mixed-effects models in the lme4 package (21: distance ~ evolutionary treatment + (1|ancestry)) to test whether each matching treatment lay significantly closer to the front than ancestral populations, using one-sided contrast tests in the emmeans package (22). Because more than one treatment can be relevant to a given relationship (e.g. both nitrogen- and phosphorus-limited populations for the 1/$N^*$-1/$P^*$ front), we considered a relationship to show significant evidence of evolutionary optimization if any matching treatment moved significantly closer to the Pareto front relative to its ancestors.

To test whether experimental evolution produced systematic changes in individual traits, we fit linear mixed-effects models using the lme4 package (21). We modeled changes in trait values (calculated relative to population-specific ancestral means) as a function of evolutionary environment (fixed effect) with ancestry included as a random effect (change in trait value ~ evolutionary treatment + (1|ancestry)). We assessed the significance of evolutionary environment using likelihood ratio tests. We quantified the contribution of ancestry to residual variance using adjusted intraclass correlation coefficients, and evaluated the relative importance of fixed versus random effects using marginal and conditional R-squared values calculated using the performance package (23). To identify the particular effects of specific evolutionary treatments on changes in *C.*

*reinhardtii* traits we estimated the marginal means of each treatment using the emmeans package (22).

The code responsible for single gradient analyses can be found in 08_intra-gradient_trade-offs.R; cross-gradient analyses are found in scripts 09_inter-gradient_trade-offs.R and 10_secondary_trait_trade-offs.R. The single-trait (univariate trait change) analysis can be found in 12_evol_effects_modelling.R.

**xii. Multivariate partitioning of phenotypic diversity and ancestral and evolutionary effects.**
To assess whether evolutionary treatment or ancestry better explained multivariate patterns of phenotypic variation, we performed redundancy analyses (RDA) using trait matrices constrained by evolutionary environment and ancestry. These analyses included all niche-determining traits, as well as pigmentation, elemental stoichiometry, and cell volume, for the 32 experimentally evolved and 5 ancestral populations of *Chlamydomonas reinhardtii*. We quantified the relative contributions of evolutionary environment and ancestry to multivariate trait variation using the rda function in the vegan package (24). To evaluate the total variance explained by each explanatory factor, we summed the canonical eigenvalues associated with the constrained axes. Because nitrogen and phosphorus content strongly dominated multivariate structure, we conducted RDAs both on the full trait set and on a reduced dataset excluding these variables (Figure S5).

We further visualized multivariate trait structure using principal component analysis (PCA) implemented in the vegan package (24). PCA was used to explore patterns of population-level differentiation and covariation among traits, rather than for formal hypothesis testing. We focused on the first two principal components (PC1 and PC2), which explained the greatest proportion of phenotypic variation, and examined trait loadings (eigenvector coefficients) to interpret how individual traits contributed to differentiation among populations and to multivariate trade-off structure (Figure S6). Together, RDA and PCA provided complementary perspectives on multivariate trait structure, allowing us to partition variance attributable to ancestry and evolutionary treatment while visualizing patterns of trait covariation and constraint.

The code responsible for performing principal component analysis and redundancy analyses can be found in the script 13_PCA_RDA.R.

**xiii. Estimation of interspecific variation in *R** and TPC traits.**
To assess the existence of trade-offs and constraints across the phytoplankton tree of life, we compiled published estimates of growth rates and niche-determining traits across light, nitrogen, phosphorus, and temperature gradients for a broad range of phytoplankton taxa spanning cyanobacteria and eukaryotic microalgae. Where raw growth data were available, we fit thermal performance and Monod curves directly to published datasets (25–29). For studies that did not report raw data, we extracted published parameter estimates (30, 31).

In fitting Lactin II TPCs to our synthesis data, Bayesian fitting approaches failed to converge reliably due to the wide variation in thermal optima and limits across taxa. We therefore fit Lactin II models using nonlinear least-squares approaches implemented in the nls.multstart and rTPC packages (3, 6). Initial models were fit with nls.multstart using broad parameter bounds to capture interspecific variation in temperature sensitivity, thermal limits, and curve shape ($\rho$: 0 – 0.5, $\lambda$: -3 – -0.001, $T_{max}$: the minimum temperature at which growth rates were quantified in a data set – the maximum temperature at which growth rates were quantified in a data set + 5°C, $\Delta T$: 0.1 – 40). We then fit more constrained models using nlsLM from the minpack.lm (32) package with starting values distributed narrowly around parameter estimates extracted from our nls.multstart models ($\rho$: max (0.001, $\rho$ - 0.05) – $\rho$ + 0.05, $\lambda$: $\lambda$ - 0.5 – min(-0.001, $\lambda$ + 0.5), $T_{max}$: max(0, $T_{max}$- 3) – $T_{max}$+ 3, $\Delta T$ : max(0.1, $\Delta T$ -2) – $\Delta T$ + 2). We calculated estimates for $T_{br}$ and $\mu_{max}$ (T) as earlier described, using 1000 bootstrapped estimates (using the boot package, 33) to generate distributions from which we extracted and reported median values. For TPC models, to control for quality fits, we excluded datasets where growth was measured at fewer than five temperatures, no growth data above the temperature at which growth was maximized in a data set (i.e. the putative $T_{opt}$), or the confidence interval around the estimate of $T_{max}$ was greater than 15% of $T_{max}$ - $T_{min}$. Because many datasets lacked sufficient low-temperature observations to estimate lower thermal limits, we imposed a conservative lower bound of minus 1.8 °C, corresponding to the freezing point of seawater, to avoid extrapolation beyond biologically plausible conditions. After visually inspecting model fits in a custom Shiny app (code available in the “Shiny-apps” folder at Zenodo, DOI: 10.5281/zenodo.19032401), we removed curves (3/228) whose model fits did not closely match the underlying data, leaving us with a total of 225 thermal performance curves across 206 taxa.

We fit Monod curves (Equation 3) to growth rate data across light, nitrogen, and phosphorus gradients using Bayesian methods as described for our experimental *Chlamydomonas reinhardtii* populations, with expanded parameter bounds for light ($k_s$: 0 – 100) and nitrogen ($k_s$: 0 – 50) to accommodate interspecific variation (see Table S10). We calculated minimum resource requirements using a mortality rate of 0.1 per day (rather than 0.56), reflecting the lower maximum growth rates typical of many phytoplankton taxa. We manually inspected model fits using custom Shiny apps for each abiotic gradient (code available in the “Shiny-apps” folder at Zenodo, DOI: 10.5281/zenodo.19032401) and excluded taxa with poor fits. After quality control, we were left with 121 curves for 110 taxa for light, 52 observations on 36 taxa for nitrogen, and 101 observations on 67 taxa for phosphorus. In comparing interspecific variation among phytoplankton with trait data from our experimentally evolved *C. reinhardtii* populations (Figure S8), we recalculated minimal resource requirements ($R^*$) for our experimental populations using the same mortality rate of 0.1.

The code responsible for fitting TPC and Monod functions to synthesized phytoplankton data can be found in the following scripts: 14_synthesis_light.R, 15_synthesis_nitrogen.R, 16_synthesis_phosphorus.R, and 17_synthesis_temperature.R.

**xiv. Testing the significance of Pareto fronts across phytoplankton.**

We applied the same Pareto front and correlational analyses used for our experimental *C. reinhardtii* populations (Supporting Methods 2.vi-x) to the synthesized phytoplankton data. We removed all taxa not resolved to the species level, and collapsed intraspecific variation to a single value for each trait for each species (N = 189) in our dataset by averaging niche-determining trait and growth rate values. For each bivariate relationship, we then assessed the significance of Pareto fronts using the empty-space randomization test and characterized trait covariation using Pearson correlations as described above. When evaluating constraints among niche-determining traits, we excluded single extreme outlier species that exceeded the 90$^{th}$ quantile on both trait axes, whose inclusion would otherwise obscure broader patterns of constraint.

The code responsible for fitting and testing Pareto fronts, and applying correlational approaches to the phytoplankton synthesis data can be found in script 18_synthesis_Pareto.R.

### Part B: Supporting Figures

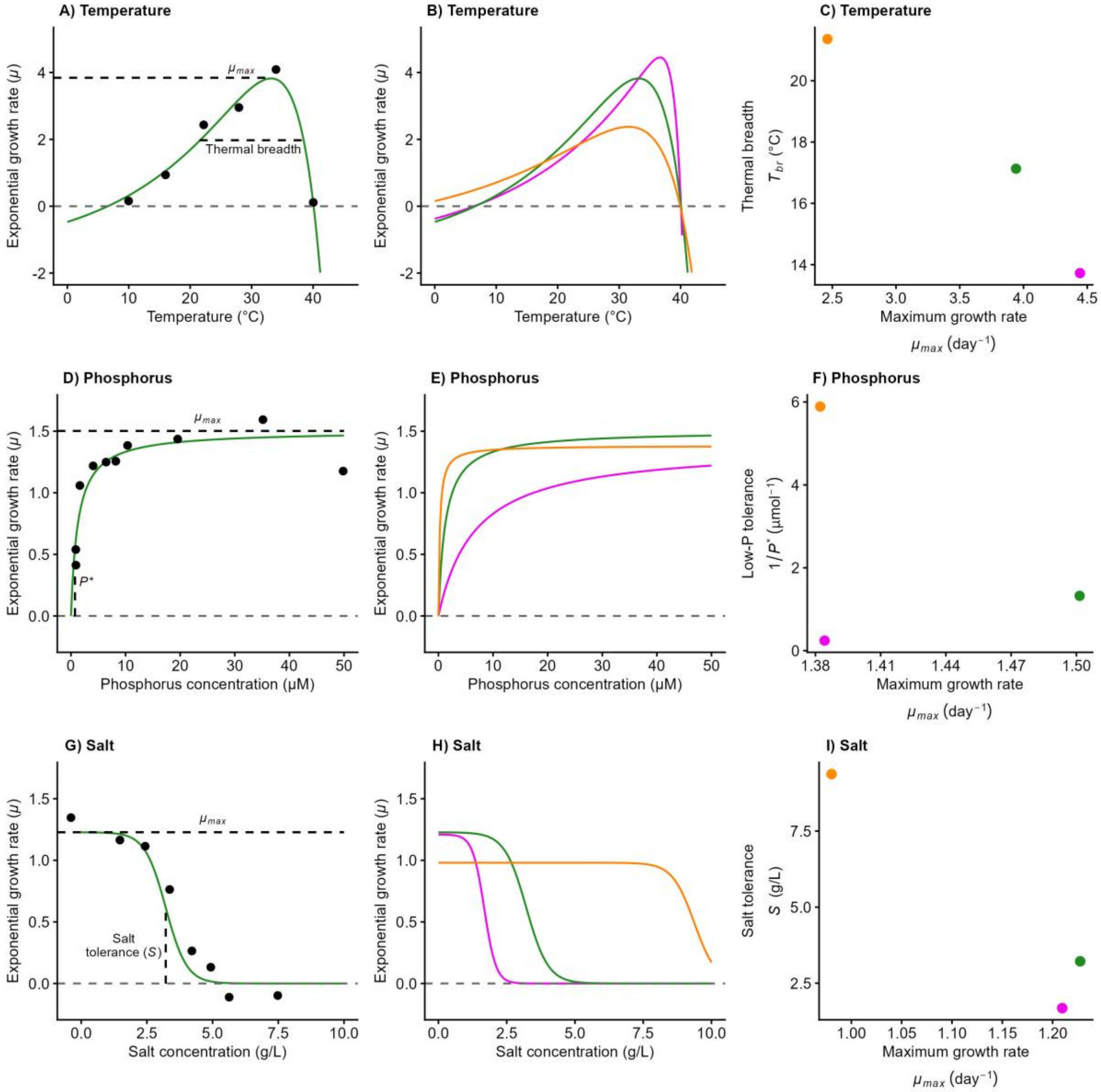

**Figure S1. Representation of model-fitting approach.**
We fit Bayesian models to the exponential growth rate ($\mu$, $day^{-1}$) of *Chlamydomonas reinhardtii* in batch culture experiments across abiotic gradients. (A) Lactin II thermal performance curve (TPC), fit to growth data from the replicate with median thermal breadth ($T_{br}$, the range of temperatures at which $\mu \geq \mu_{max}/2$). Dashed lines identify maximum exponential growth rate ($\mu_{max}$) and thermal breadth. (B) Lactin II TPCs for replicates with median (green), minimum (purple), and maximum (orange) $T_{br}$. Corresponding estimates of $T_{br}$ and $\mu_{max}$ plotted in (C). (D) Monod curve fitting variation in $\mu$ across a nutrient gradient (here phosphorus) for the replicate with median $1/P^*$ (low-phosphorus tolerance). Dashed lines highlight $\mu_{max}$ and $R^*$ (here $P^*$), the minimum resource requirement for population growth. (E) Monod curves for replicates with minimum, maximum, and median $1/P^*$ values, for which $\mu_{max}$ and $1/P^*$ are represented in (F). (G) Inverse logistic growth curve modeling exponential growth rates of the population with median salt tolerance (*S*) across a salinity gradient. Dashed lines indicate $\mu_{max}$ and *S*, the concentration of salt at which $\mu \leq \mu_{max}/2$. (H) Salt tolerance curves for populations with minimum, maximum, and median salt tolerance, for which *S* and $\mu_{max}$ are plotted in (I).

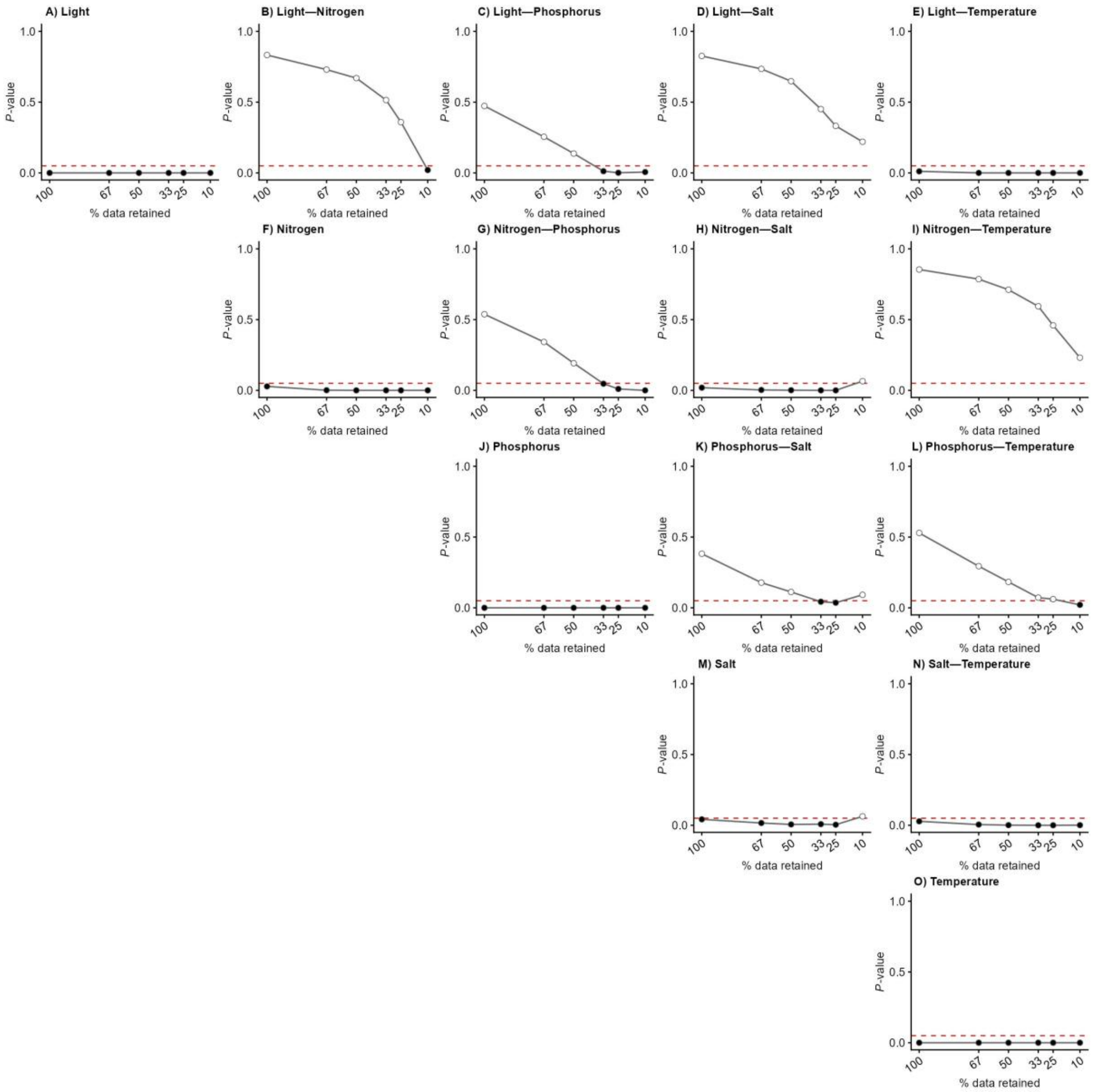

**Figure S2. Sensitivity of Pareto front significance to variation in data retention thresholds.** Each panel shows the *P* value from the Pareto front analysis (empty-space randomization test) as data are progressively excluded on the basis of scaled Euclidean distance from the minimum (x, y) point. Dashed red lines indicate $P$ = 0.05; filled circles represent significant results; open circles represent non-significant results. Diagonal panels (A, F, J, M, O) show results for relationships between $\mu_{max}$ and niche-determining traits ($1/R^*$, $S$, and $T_{br}$) within single abiotic gradients. Off-diagonal panels show results for pairwise comparisons between niche-determining traits across niche axes.

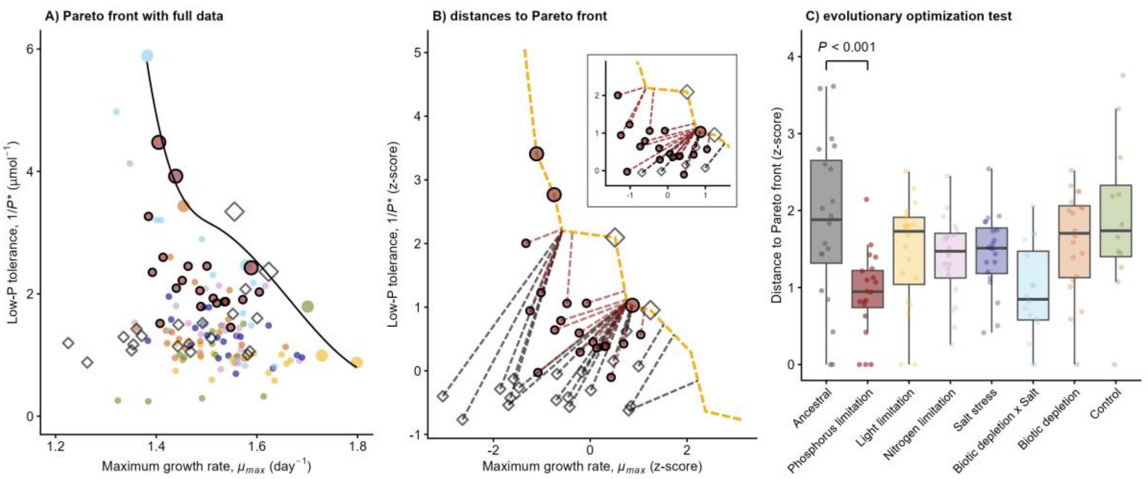

**Figure S3. Schematic representation of evolutionary optimization methods and inference.** (A) As in Figures 2-4 of the main text, a monotonic, decreasing smooth Pareto front is fit to Pareto optimal points (larger points) using the 'scam' package. All data are shown, with populations whose experimental evolutionary conditions match those of the panel-specific abiotic gradients (here phosphorus) outlined in black. Ancestral populations' trait values are shown as diamonds. (B) Depiction of the corresponding stepwise (un-smoothed) Pareto front used for statistical testing (shown in dashed yellow). After z-scoring each trait, we calculated each replicate's Euclidean distance to the nearest point on the Pareto front (shown as dashed lines connecting ancestral (in diamonds) and descendant phosphorus-limited populations (in circles) to the Pareto front). Inset: equal x- and y-axis ranges confirm that dashed lines are perpendicular to the Pareto front, validating closest-point assignments. (C) Boxplot depicting the distance from the Pareto front for ancestral and experimentally evolved *C. reinhardtii* populations. We report the $P$ value from a one-sided contrast in our mixed-effects model, testing whether matching (here phosphorus-limited) populations lie closer to the Pareto front than ancestral populations.

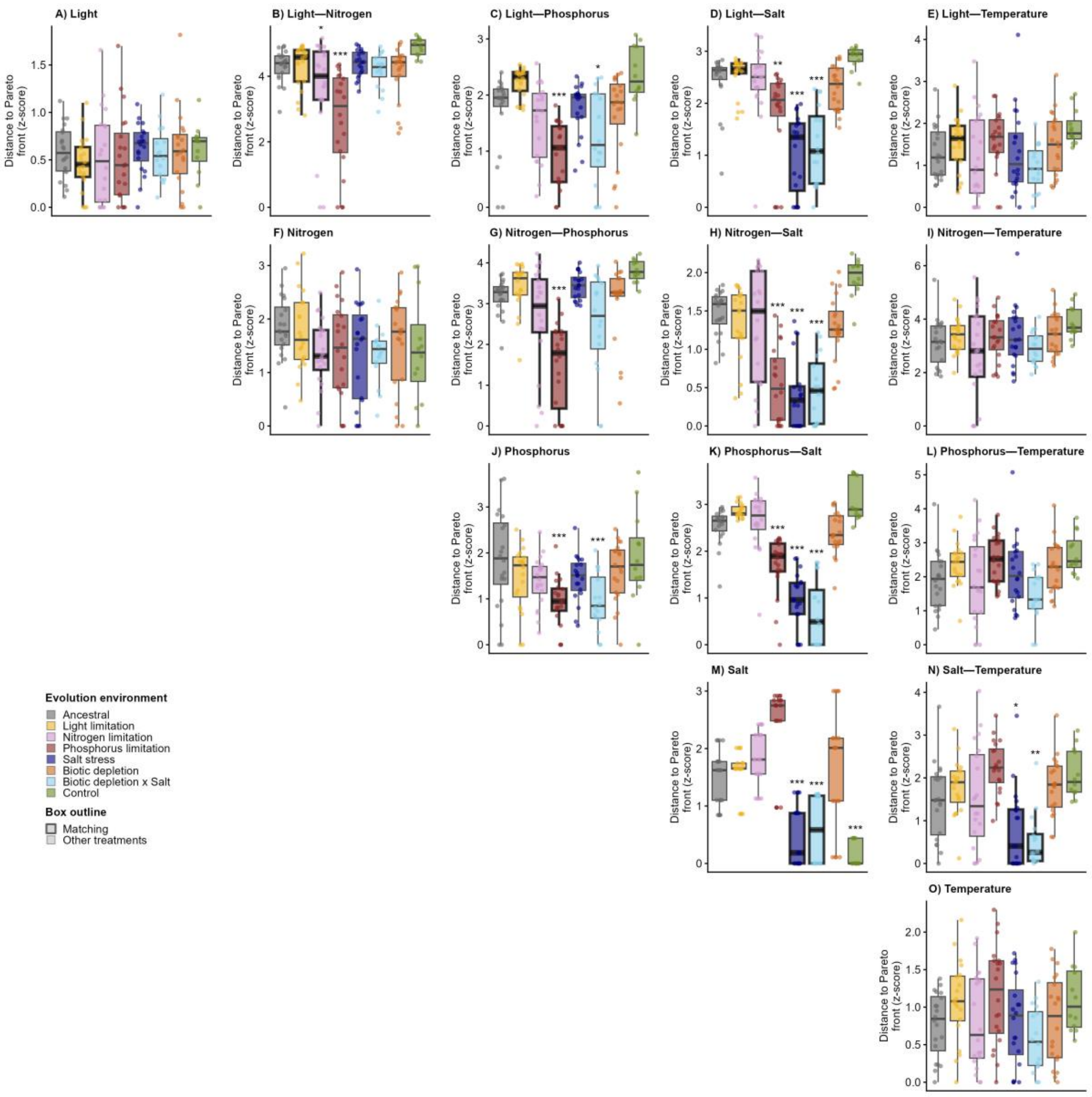

**Figure S4. Evolutionary optimization of niche-determining traits across niche axes in *C. reinhardtii*.**

Each panel (A–O) shows the scaled (z-scored) Euclidean distance of ancestral and experimentally evolved populations from the Pareto front for one bivariate trait relationship; smaller distances indicate trait combinations closer to the front (i.e. greater optimization). Boxes are coloured by evolution environment (ancestral; light-, nitrogen-, and phosphorus-limited; salt-stressed; biotically depleted; biotic depletion × salt; and control) and outlined in black for treatments whose selection environment matches the panel's niche axis (or axes). Diagonal panels (A, F, J, M, O) show relationships between $\mu_{max}$ and single niche-determining traits within individual gradients; off-diagonal panels show relationships between niche-determining traits across gradients. For each treatment, we used a one-sided contrast in a mixed-effects model (distance ~ selection environment + (1 | ancestry)) to test whether its populations lay significantly closer to the Pareto front than ancestral populations; asterisks denote significance (* $P < 0.05$, ** $P < 0.01$, *** $P < 0.001$), with full statistics in Table S3.

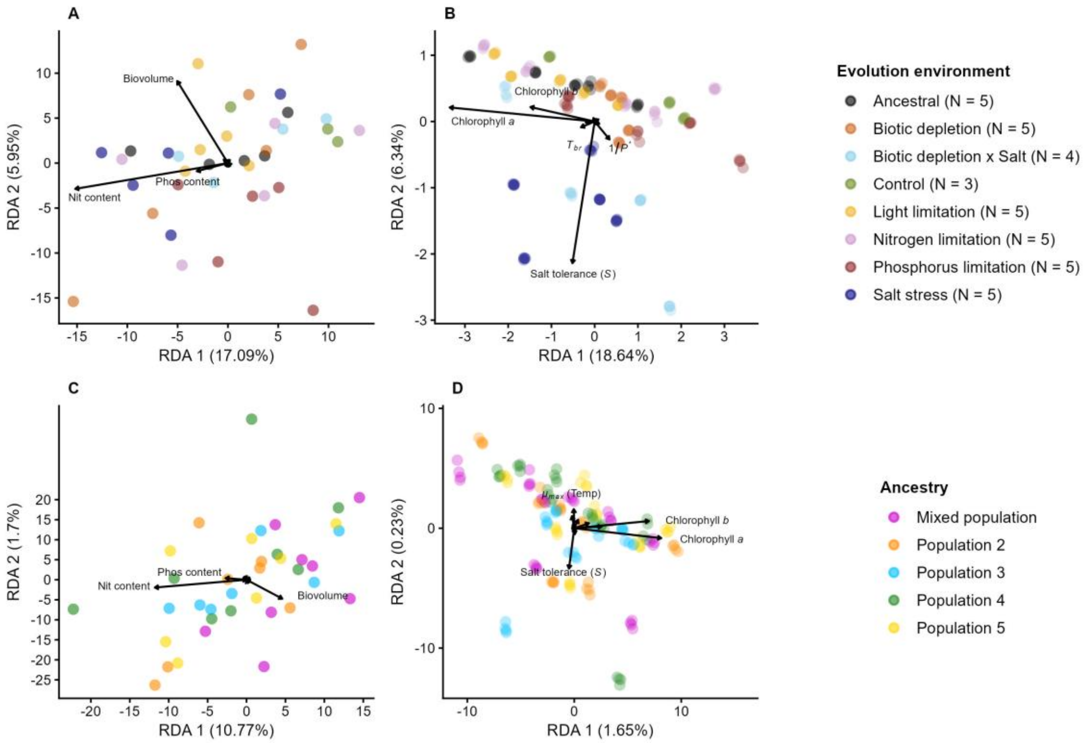

**Figure S5. Redundancy analyses of *C. reinhardtii* traits.**
We performed redundancy analyses of cell morphology, pigmentation, and stoichiometry alongside niche-determining traits, of experimentally evolved and ancestral *Chlamydomonas reinhardtii* replicates, constrained by (A-B) selection environment and (C-D) ancestry. Points represent individual replicates, coloured by evolution environment (A–B) or ancestral population (C–D); arrows show trait loadings on the first two RDA axes. (A) Evolution environment explains 23.18% of total variation, with RDA axes 1 and 2 accounting for 73.73% and 25.67% of the variation explained by evolutionary history. (B) After removing stoichiometric data and biovolume, the evolution environment explains 25.86% of total variation, with RDA axes 1 and 2 capturing 72.08% and 24.52% of the variation explained by evolutionary history. (C) Ancestry explains 12.53% of total variation, with RDA axes 1 and 2 accounting for 85.95% and 13.57% of the variation accounted for by ancestry. (D) After removing biovolume and stoichiometric data, ancestry accounts for only 2.06% of total variation, with RDA axes 1 and 2 explaining 80.10% and 11.17% of the variation explained by ancestry.

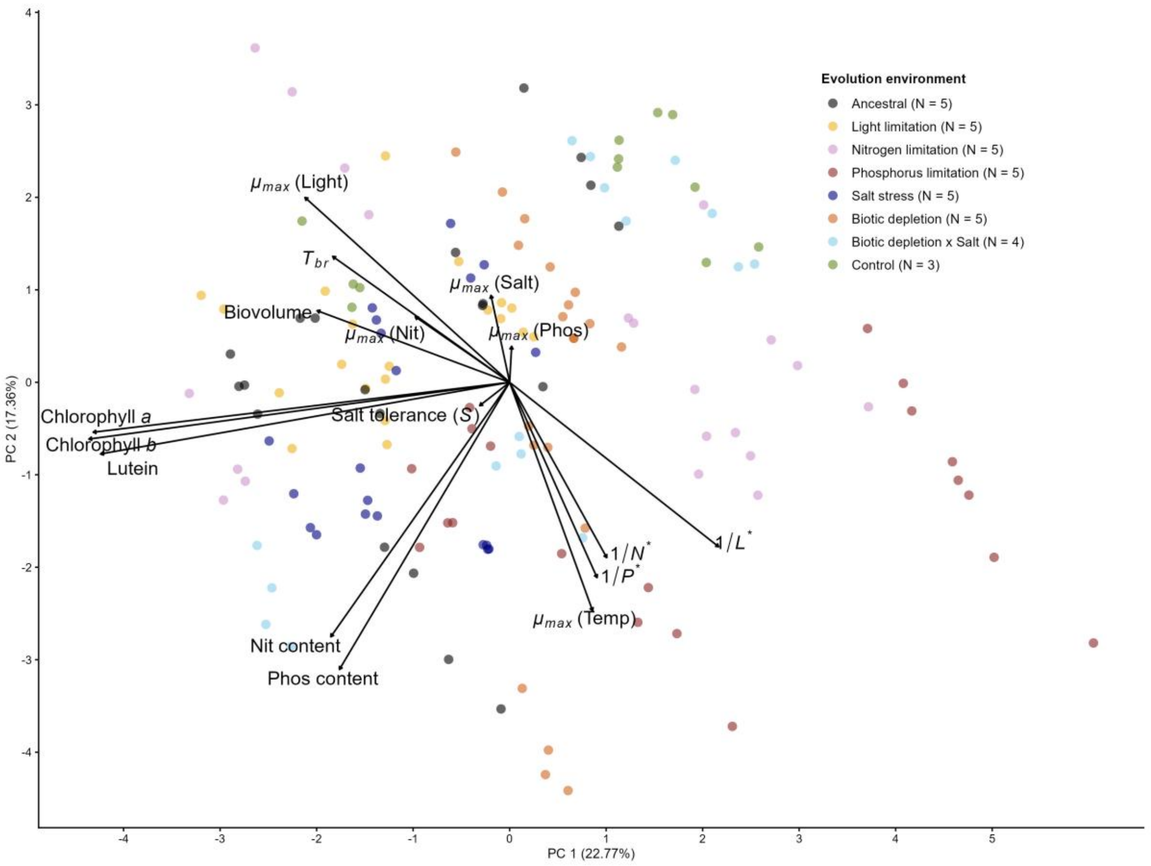

**Figure S6. Principal Component Analysis (PCA) of *C. reinhardtii* traits.**
We measured cell morphology, pigmentation, and stoichiometry alongside niche-determining traits across experimentally evolved and ancestral *Chlamydomonas reinhardtii* replicates. Points represent individual replicates, coloured by evolution environment; arrows show the loadings of each trait on PC 1 and PC 2, with opposing arrows indicating negative correlations between traits. Together, PC 1 and PC 2 explain 40.13% of total variation. Each evolutionary treatment comprised five populations, except the control (N = 3) and biotic depletion × salt (N = 4). Each point represents trait data from one of four replicate assays per population (salt, stoichiometric, biovolume, and pigmentation assays were not replicated).

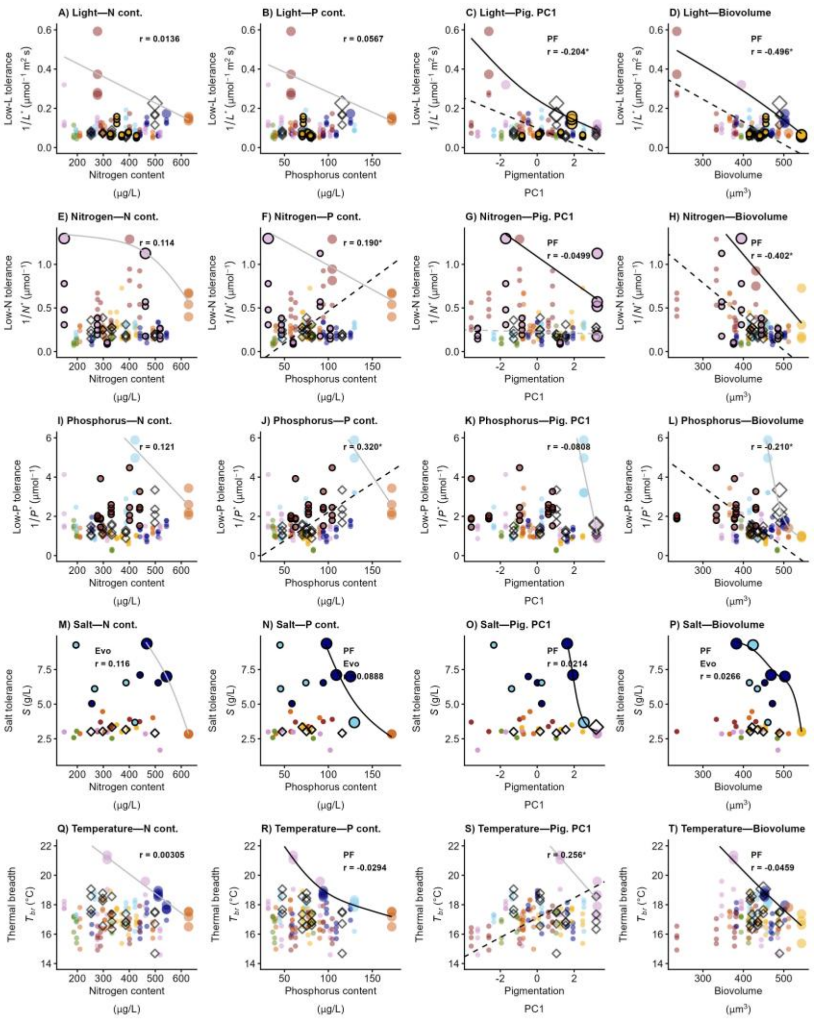

**Figure S7. Trade-offs between niche-determining traits, stoichiometry, pigmentation, and biovolume.**

Trade-offs between low-resource tolerance (1/$R^*$) for light, nitrogen, phosphorus, thermal breadth ($T_{br}$) and salt tolerance ($S$) with stoichiometric data (nitrogen and phosphorus content (cont.)), pigmentation (Pig. PC1), and biovolume in experimentally evolved and ancestral *Chlamydomonas reinhardtii* populations. Because the three pigmentation variables (lutein, chlorophyll *a*, and chlorophyll *b*) were highly collinear (~96% of their variance is captured by PC1 in a pigment-exclusive PCA), we summarize pigmentation here using the PC1 scores from this analysis. Trait data for populations that evolved under the relevant selective environment(s) (e.g. under light limitation in (A)) are shown as filled circles outlined in black; other experimentally evolved populations are filled circles without outlines. Ancestral populations are represented as open diamonds. Each point represents trait data from one of four replicate assays per population (salt, stoichiometric, biovolume, and pigmentation assays were not replicated). Smoothed black lines—representing putative Pareto fronts—were fit to the set of optimal trait combinations (identified via a convex hull algorithm) using smoothed, monotonically decreasing curves. Pareto-optimal trait combinations are shown as larger circles and diamonds. Dashed lines are standardized major axis (SMA) regressions, plotted only where the Pearson correlation between traits was significant (*r* reported in each panel; asterisk denotes significance at $\alpha$ = 0.05). Significant Pareto fronts (which define larger inaccessible spaces than null simulations) are plotted in black; non-significant Pareto fronts are shown in grey. Text labels report the correlation (*r*) and significant results ($\alpha$ = 0.05; PF: a significant Pareto front constrains phenotypic variation; Evo: experimental evolution has moved at least one matching population significantly closer to the Pareto front). All data are plotted.

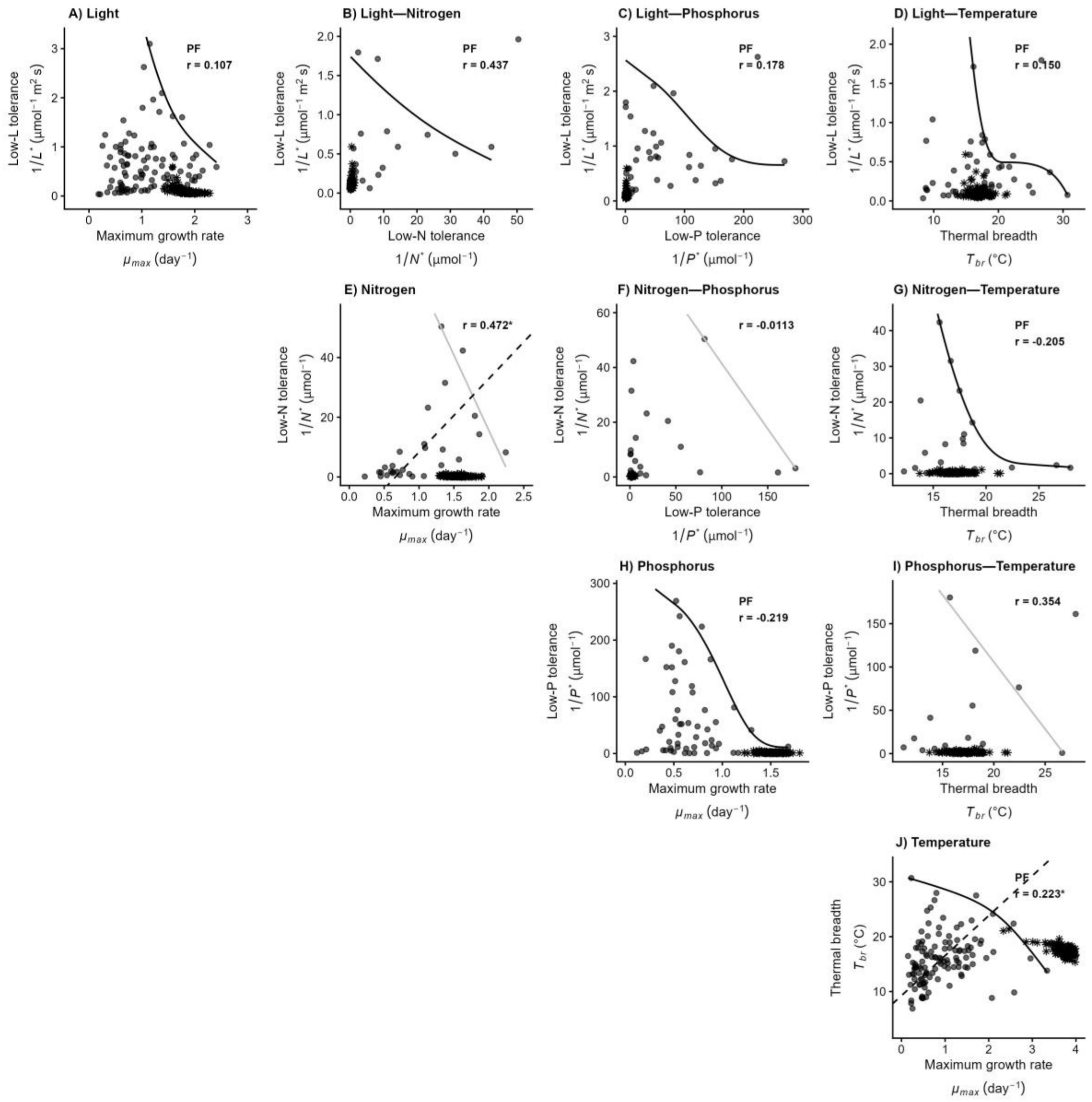

**Figure S8. Trade-offs across phytoplankton with our *C. reinhardtii* data added.**
Trade-offs between maximum specific growth rates ($\mu_{max}$), low-resource tolerance ($1/R^*$) for light, nitrogen, and phosphorus, and thermal breadth ($T_{br}$) across 189 phytoplankton species. All circular data points represent the mean trait values of a single phytoplankton species; apparent variation in point shading reflects the plotting of overlapping values. Data from our experimentally evolved *Chlamydomonas reinhardtii* populations are shown as black asterisk markers. Solid lines represent bivariate Pareto fronts constraining trait co-optimization using smoothed, monotonically decreasing curves. Dashed lines are standardized major axis (SMA) regressions, plotted only where the Pearson correlation between traits was significant (r reported in each panel; asterisk denotes significance at $\alpha = 0.05$). Text labels report the correlation (r) and significant results ($\alpha = 0.05$; PF: a significant Pareto front constrains phenotypic variation). Significant Pareto fronts are shown in black; non-significant fronts are light grey.

### Part C: Supporting Tables

**Table S1. Sensitivity of Pareto front significance to variation in data retention thresholds.** To test for significant Pareto fronts, we first removed sub-optimal data points prior to permutation testing. This step was necessary because a large proportion of our data derive from populations that were not under strong selection to optimize the trait relationships in question—including data points in the interior of trait space that are uninformative about upper boundary structure. We report permutation *P* values across six data retention thresholds: 100% (all data), 66.67%, 50%, 33.33%, 25%, and 10%, where retention is based on scaled Euclidean distance from the sub-optimal corner (minimum x,y values) of trait space. Bold *P* values indicate significant Pareto fronts (≤ 0.05). All results reported in the main text use the 33.33% threshold, which retains only the upper third of data points and was selected as a balance between statistical power and focus on the performance frontier. Each cell reports the *P* value for the relationship between the row and column gradients at each retention threshold; diagonal cells (e.g. Light × Light) are within-axis relationships ($\mu_{max}$ vs. the corresponding trait), and off-diagonal cells are cross-axis relationships between two niche-determining traits.

| | | **Gradient** | | | | |
|---|---|---|---|---|---|---|
| **Data threshold (%)** | **Gradient** | **Light** | **Nitrogen** | **Phosphorus** | **Salt** | **Temperature** |
| **100 (all)** | **Light** | **<0.001** | 0.833 | 0.473 | 0.826 | **0.011** |
| **66.67** | | **<0.001** | 0.73 | 0.255 | 0.735 | **<0.001** |
| **50** | | **<0.001** | 0.67 | 0.137 | 0.648 | **<0.001** |
| **33.33** | | **<0.001** | 0.515 | **0.012** | 0.451 | **<0.001** |
| **25** | | **<0.001** | 0.359 | **0.001** | 0.332 | **<0.001** |
| **10** | | **<0.001** | **0.02** | **0.006** | 0.220 | **<0.001** |
| **100 (all)** | **Nitrogen** | | **0.028** | 0.538 | **0.019** | 0.854 |
| **66.67** | | | **0.001** | 0.342 | **0.003** | 0.786 |
| **50** | | | **<0.001** | 0.192 | **0.001** | 0.711 |
| **33.33** | | | **<0.001** | **0.047** | **<0.001** | 0.594 |
| **25** | | | **<0.001** | **0.01** | **<0.001** | 0.459 |
| **10** | | | **<0.001** | **<0.001** | 0.065 | 0.231 |
| **100 (all)** | **Phosphorus** | | | **<0.001** | 0.382 | 0.529 |
| **66.67** | | | | **<0.001** | 0.178 | 0.294 |
| **50** | | | | **<0.001** | 0.112 | 0.183 |
| **33.33** | | | | **<0.001** | **0.043** | 0.072 |
| **25** | | | | **<0.001** | **0.036** | 0.061 |
| **10** | | | | **<0.001** | 0.092 | **0.021** |
| **100 (all)** | **Salt** | | | | **0.042** | **0.028** |
| **66.67** | | | | | **0.016** | **0.005** |
| **50** | | | | | **0.006** | **0.001** |
| **33.33** | | | | | **0.008** | **<0.001** |
| **25** | | | | | **0.004** | **<0.001** |
| **10** | | | | | 0.063 | **0.001** |
| **100 (all)** | **Temperature** | | | | | **<0.001** |
| **66.67** | | | | | | **<0.001** |
| **50** | | | | | | **<0.001** |
| **33.33** | | | | | | **<0.001** |
| **25** | | | | | | **<0.001** |
| **10** | | | | | | **<0.001** |

**Table S2. Significance of Pareto fronts and evolutionary optimization in *C. reinhardtii*.** Diagonal cells show within-axis relationships between $\mu_{max}$ and niche-determining traits (1/$R^*$, $T_{br}$, salt tolerance); off-diagonal cells show relationships among niche-determining traits across axes. For each relationship we report the empty-space randomization test for a Pareto front (Li et al. (33%)), the Pearson correlation (*r*, with its *P* value in brackets), and a test of evolutionary optimization (Evol; mixed-effects model of distance to the front, reported per matching treatment by single-letter code; Methods). Values are n/a for evolutionary comparisons involving only thermal traits (no treatment imposed thermal stress); Evol = 1* indicates ancestral populations lie closer to the front than matching populations. Significant *P* values (≤ 0.05) are bolded.

| | | **Gradient** | | | | |
|---|---|---|---|---|---|---|
| **Statistical test** | **Gradient** | **Light (L)** | **Nitrogen (N)** | **Phosphorus (P)** | **Salt (Salt: S, Salt x Biotic depletion: BS)** | **Temperature (T)** |
| **Li et al. (33%)<br>Pearson's r<br>Evol** | **Light (L)** | **<0.001**<br>**-0.503 (<0.001)**<br>L: 0.676 | 0.515<br>**0.313 (<0.001)**<br>L: 0.991;<br>**N: 0.0436** | **0.012**<br>**0.237 (0.00373)**<br>L: 1*; **P: <0.001** | 0.451<br>0.00973 (0.906)<br>L: 1*; **S: <0.001;**<br>**BS: <0.001** | **<0.001**<br>-0.159 (0.0535)<br>L: 1* |
| **Li et al. (33%)<br>Pearson's r<br>Evol** | **Nitrogen (N)** | | **<0.001**<br>**-0.317 (<0.001)**<br>N: 0.196 | **0.047**<br>**0.284 (<0.001)**<br>N: 0.154;<br>**P: <0.001** | **<0.001**<br>**-0.167 (0.0419)**<br>N: 0.620;<br>**S: <0.001;**<br>**BS: <0.001** | 0.594<br>-0.0291 (0.725)<br>N: 0.440 |
| **Li et al. (33%)<br>Pearson's r<br>Evol** | **Phosphorus (P)** | | | **<0.001**<br>**-0.289 (<0.001)**<br>**P: <0.001** | **0.043**<br>-0.0217 (0.794)<br>**P: <0.001;**<br>**S: <0.001;**<br>**BS: <0.001** | 0.072<br>-0.0600 (0.468)<br>P: 1* |
| **Li et al. (33%)<br>Pearson's r<br>Evol** | **Salt (Salt: S, Salt x Biotic depletion: BS)** | | | | **0.008**<br>**-0.175 (0.0332)**<br>**S: <0.001;**<br>**BS: <0.001** | **<0.001**<br>0.134 (0.105)<br>**S: 0.0244;**<br>**BS: 0.0015** |
| **Li et al. (33%)<br>Pearson's r<br>Evol** | **Temperature (T)** | | | | | **<0.001**<br>**-0.664 (<0.001)**<br>n/a |

**Table S3. Evolutionary treatment effects on distance to Pareto fronts in *C. reinhardtii***
For each bivariate trait relationship (a Pareto front defined by a pair of niche axes) and each evolutionary treatment, we report the treatment's effect on populations' scaled Euclidean distance to the front relative to ancestral populations, estimated from a mixed-effects model (distance ~ evolutionary history + (1|ancestry)). Row blocks give the first niche axis of each relationship and columns the second; sub-rows within each block give the evolutionary treatment. Each cell shows the estimated change in distance with its *P* value in brackets; negative estimates indicate movement toward the front (i.e. greater optimization). This table reports the full per-treatment results underlying the Evol summaries in Table S2. Significant *P* values (≤ 0.05) are bolded.

| | | Gradient | | | | |
|---|---|---|---|---|---|---|
| **Evolutionary Treatment** | **Gradient** | **Light** | **Nitrogen** | **Phosphorus** | **Salt (S)** | **Temperature** |
| **Light** | **Light** | -0.122 (0.676) | -0.00961 (0.991) | 0.515 (1) | 0.106 (1) | 0.241 (1) |
| **Nitrogen** | | -0.0586 (0.924) | **-0.721 (0.0436)** | -0.164 (0.742) | -0.00213 (0.992) | -0.132 (0.902) |
| **Phosphorus** | | -0.0580 (0.925) | **-1.677 (<0.001)** | **-0.749 (<0.001)** | **-0.688 (0.0018)** | 0.284 (1) |
| **Salt** | | 0.0443 (0.999) | 0.06449 (0.998) | 0.0537 (0.999) | **-1.391 (<0.001)** | 0.0151 (0.995) |
| **Salt X Biotic depletion** | | -0.0146 (0.985) | -0.160 (0.918) | **-0.496 (0.0319)** | **-1.267 (<0.001)** | -0.444 (0.228) |
| **Biotic depletion** | | 0.0220 (0.998) | -0.209 (0.843) | -0.0511 (0.967) | -0.131 (0.866) | 0.192 (1) |
| **Control** | | 0.0263 (0.998) | 0.722 (1) | 0.726 (1) | 0.525 (1) | 0.743 (1) |
| **Light** | **Nitrogen** | | -0.065 (0.971) | 0.223 (1) | -0.132 (0.773) | 0.235 (1) |
| **Nitrogen** | | | -0.466 (0.196) | -0.539 (0.154) | -0.170 (0.620) | -0.445 (0.440) |
| **Phosphorus** | | | -0.425 (0.270) | **-1.680 (<0.001)** | **-0.937 (<0.001)** | 0.164 (1) |
| **Salt** | | | -0.431 (0.260) | 0.266 (1) | **-1.049 (<0.001)** | 0.226 (1) |
| **Salt X Biotic depletion** | | | -0.490 (0.209) | -0.689 (0.0619) | **-1.007 (<0.001)** | -0.262 (0.822) |
| **Biotic depletion** | | | -0.275 (0.636) | -0.162 (0.894) | -0.198 (0.500) | 0.282 (1) |
| **Control** | | | -0.387 (0.494) | 0.690 (1) | 0.478 (1) | 0.744 (1) |
| **Light** | **Phosphorus** | | | -0.396 (0.228) | 0.327 (1) | 0.475 (1) |
| **Nitrogen** | | | | -0.464 (0.120) | 0.112 (1) | -0.0436 (0.982) |
| **Phosphorus** | | | | **-0.927 (<0.001)** | **-0.834 (<0.001)** | 0.593 (1) |
| **Salt** | | | | -0.401 (0.219) | **-1.587 (<0.001)** | 0.232 (1) |
| **Salt X Biotic depletion** | | | | **-0.938 (<0.001)** | **-1.876 (<0.001)** | -0.541 (0.204) |
| **Biotic depletion** | | | | -0.258 (0.593) | -0.152 (0.743) | 0.383 (1) |
| **Control** | | | | 0.153 (1) | 0.537 (1) | 0.860 (1) |
| **Light** | **Salt** | | | | 0.0864 (1) | 0.316 (1) |
| **Nitrogen** | | | | | 0.334 (1) | 0.0679 (0.999) |
| **Phosphorus** | | | | | 0.896 (1) | 0.778 (1) |
| **Salt** | | | | | **-1.037 (<0.001)** | **-0.698 (0.0244)** |
| **Salt X Biotic depletion** | | | | | **-0.941 (<0.001)** | **-0.984 (0.0015)** |
| **Biotic depletion** | | | | | 0.182 (1) | 0.342 (1) |
| **Control** | | | | | **-1.219 (<0.001)** | 0.661 (1) |
| **Light** | **Temperature** | | | | | 0.306 (1) |
| **Nitrogen** | | | | | | 0.0544 (0.999) |
| **Phosphorus** | | | | | | 0.395 (1) |
| **Salt** | | | | | | 0.0488 (0.999) |
| **Salt X Biotic depletion** | | | | | | -0.288 (0.290) |
| **Biotic depletion** | | | | | | 0.0642 (0.999) |
| **Control** | | | | | | 0.450 (1) |

**Table S4. Linear mixed model results.**

We used linear mixed effects models (Δtrait ~ Evolutionary treatment + (1|Ancestry)) to test whether experimental evolution shifted trait values, and if so, in which direction. For each trait we report the significance of the evolution effect (likelihood-ratio test), the proportion of variance attributable to shared ancestry (adjusted ICC), the conditional and marginal $R^2$, and the estimated marginal mean change under each selection environment. Significant values ($P \leq 0.05$) are bolded.

| Change in trait | Evolution (LRT *P* value) | Ancestry (proportion of variation) | $R^2$ (conditional) | $R^2$ (marginal) | Biotic depletion | Biotic x salt stress | Control | Light limitation | Nitrogen limitation | Phosphorus limitation | Salt stress |
|---|---|---|---|---|---|---|---|---|---|---|---|
| $1/L^*$ | **<0.001** | 0.398 | 0.474 | 0.125 | 0.00570 | 0.0153 | -0.0334 | -0.00611 | 0.0218 | **0.0680** | 0.0105 |
| $\mu_{max}$ (L) | **<0.001** | 0.673 | 0.695 | 0.068 | -0.0546 | -0.0783 | 0.1001 | 0.0696 | -0.0469 | -0.1100 | -0.0818 |
| $1/N^*$ | **<0.001** | 0.054 | 0.332 | 0.294 | 0.0822 | 0.0250 | -0.0742 | 0.0576 | **0.173** | **0.343** | -0.0200 |
| $\mu_{max}$ (N) | 0.0622 | 0.276 | 0.326 | 0.068 | 0.000908 | 0.0459 | 0.0792 | -0.0154 | 0.00721 | -0.0292 | 0.0686 |
| $1/P^*$ | **<0.001** | 0.649 | 0.706 | 0.162 | -0.209 | 0.598 | -0.750 | -0.743 | -0.160 | 0.650 | -0.389 |
| $\mu_{max}$ (P) | **0.0172** | 0.544 | 0.569 | 0.056 | 0.0475 | 0.0690 | 0.0535 | **0.132** | 0.0796 | 0.0452 | 0.0910 |
| Salt tolerance (*S*) | **<0.001** | 0.040 | 0.746 | 0.735 | 0.348 | **3.310** | -0.290 | 0.153 | -0.365 | 0.461 | **3.923** |
| $\mu_{max}$ (S) | **<0.001** | 0.323 | 0.491 | 0.249 | -0.0645 | 0.0559 | **0.269** | -0.0176 | -0.0603 | **-0.178** | -0.0136 |
| $T_{br}$ | **<0.001** | 0.635 | 0.661 | 0.073 | -0.560 | 0.333 | -0.922 | -0.467 | 0.0829 | -1.126 | -0.307 |
| $\mu_{max}$ (T) | **0.0147** | 0.693 | 0.705 | 0.039 | 0.324 | 0.0799 | 0.174 | 0.0666 | -0.0669 | 0.323 | 0.164 |
| N content | **<0.001** | 0.426 | 0.508 | 0.143 | 32.67 | -33.91 | -105.49 | 21.02 | -9.41 | -6.01 | 90.01 |
| P content | **<0.001** | 0.430 | 0.489 | 0.104 | 5.70 | -4.81 | -24.55 | -2.49 | -11.80 | 1.23 | 19.04 |
| Pigmentation | **<0.001** | 0.605 | 0.643 | 0.096 | -1.504 | -0.982 | -1.901 | 0.0426 | -1.183 | -2.074 | -0.264 |
| Biovolume | **<0.001** | 0.391 | 0.584 | 0.316 | -2.52 | -11.67 | -7.88 | 19.39 | -33.24 | **-90.76** | 0.509 |

**Table S5. Significance of Pareto constraints across phytoplankton.** To characterize multivariate phenotypic constraints across light, nitrogen, phosphorus, and temperature gradients, we synthesized niche-determining trait data for 189 phytoplankton species. We tested the significance of putative Pareto fronts constraining bivariate trait optimization by comparing the observed inaccessible space to null randomizations, using the top third of phenotypic data furthest from the minimal (*x*, *y*) values by scaled Euclidean distance (Li et al. (33%)). For each relationship we also report the Pearson correlation (*r*) with its *P* value in brackets. Diagonal cells show within-axis relationships between $\mu_{max}$ and niche-determining traits; off-diagonal cells show relationships among niche-determining traits across axes. Significant *P* values (≤ 0.05) are bolded.

| | | **Gradient** | | | |
|---|---|---|---|---|---|
| **Statistical test** | **Gradient** | **Light** | **Nitrogen** | **Phosphorus** | **Temperature** |
| **Li et al. (33%)**<br>**Pearson's r** | **Light** | **0.002**<br>0.107 (0.294) | **0.002**<br>0.437 (0.0540) | **0.010**<br>0.178 (0.286) | **0.005**<br>0.150 (0.321) |
| **Li et al. (33%)**<br>**Pearson's r** | **Nitrogen** | | 0.054<br>**0.472 (0.00844)** | 0.288<br>-0.0113 (0.960) | **<0.001**<br>-0.205 (0.373) |
| **Li et al. (33%)**<br>**Pearson's r** | **Phosphorus** | | | **<0.001**<br>-0.219 (0.101) | 0.127<br>0.354 (0.106) |
| **Li et al. (33%)**<br>**Pearson's r** | **Temperature** | | | | **0.022**<br>**0.223 (0.0158)** |

**Table S6. Significance of Pareto fronts and evolutionary optimization with additional traits in *C. reinhardtii*.**

We tested relationships between niche-determining traits (1/$R^*$, $T_{br}$, salt tolerance) and stoichiometric (N and P content), pigmentation, and biovolume data; pigmentation is summarized as PC1 of a pigment-exclusive PCA (~96% of variance). For each relationship we report the empty-space randomization test for a Pareto front on the top third of data (Li et al. (33%)), the Pearson correlation (*r*, with *P* value in brackets), and an evolutionary optimization test (Evol) from a mixed-effects model of each population's distance to the front, reported per matching treatment. Values are n/a for evolutionary comparisons involving only thermal traits (no treatment imposed thermal stress); Evol = 1* indicates ancestral populations lie closer to the front than matching populations. Significant *P* values (≤ 0.05) are bolded.

| | | **Trait** | | | |
|---|---|---|---|---|---|
| **Statistical test** | **Gradient** | **N content** | **P content** | **Pigmentation** | **Biovolume** |
| **Li et al. (33%)** | **Light** | 0.660 | 0.731 | **0.007** | **0.032** |
| **Pearson's r** | | 0.0136 (0.869) | 0.0567 (0.494) | **-0.204 (0.0129)** | **-0.496 (<0.001)** |
| **Evol** | | L: 0.991 | L: 0.812 | L: 1* | L: 0.732 |
| **Li et al. (33%)** | **Nitrogen** | 0.235 | 0.102 | **0.005** | **0.004** |
| **Pearson's r** | | 0.114 (0.169) | **0.190 (0.0204)** | -0.0499 (0.547) | **-0.402 (<0.001)** |
| **Evol** | | N: 0.553 | N: 0.792 | N: 1* | N: 1* |
| **Li et al. (33%)** | **Phosphorus** | 0.655 | 0.896 | 0.807 | 0.068 |
| **Pearson's r** | | 0.1121 (0.142) | **0.320 (<0.001)** | -0.0808 (0.329) | **-0.210 (0.0102)** |
| **Evol** | | P: 0.517 | P: 0.548 | P: 1* | P: 1* |
| **Li et al. (33%)** | **Salt** | 0.132 | **0.002** | **0.004** | **0.021** |
| **Pearson's r** | | 0.116 (0.159) | 0.0888 (0.283) | 0.0214 (0.797) | 0.0266 (0.748) |
| **Evol** | | **S: <0.001;** BS: 0.477 | **S: <0.001;** BS: 0.125 | S: 0.5834; BS: 1* | **S: <0.001; BS: <0.001** |
| **Li et al. (33%)** | **Temperature** | 0.068 | **0.016** | 0.319 | **0.002** |
| **Pearson's r** | | 0.00305 (0.971) | -0.0294 (0.723) | **0.256 (0.00165)** | -0.0459 (0.579) |
| **Evol** | | n/a | n/a | n/a | n/a |

**Table S7. Design details for the experimental evolution of *C. reinhardtii*.**
We ramped up the severity of experimental stressors month by month: resource concentrations (nitrogen, phosphorus, light) were progressively reduced, while salt and biotic-depletion levels were progressively increased.

| Month | Nitrogen (N) (µM N) | Phosphorus (P) (µM P) | Light (L) (µM photons $m^{-2}s^{-1}$) | Salt (S) (g/L NaCl) | Biotically-depleted media (B) (proportion) | Biotically-depleted x Salt (BS) (proportion, g/L NaCl) | |
|---|---|---|---|---|---|---|---|
| 1 | 1000 | 50 | 100 | 0 | 0 | 0 | 0 |
| 2 | 100 | 5 | 70 | 1 | 0.01 | 0.01 | 1 |
| 3 | 100 | 5 | 50 | 2 | 0.1 | 0.1 | 2 |
| 4 | 10 | 0.5 | 20 | 4 | 0.4 | 0.4 | 4 |
| 5 | 10 | 0.5 | 15 | 6 | 0.75 | 0.75 | 6 |
| 6 | 10 | 0.5 | 5 | 8 | 0.95 | 0.95 | 8 |
| 7 | 10 | 0.5 | 5 | 8 | 1 | 1 | 8 |
| 8+ | 10 | 0.5 | 5 | 8 | 1 | 1 | 8 |

**Table S8. Specification of gradient concentrations for batch culture experiments.** We conducted growth assays across niche axes to determine the effects of experimental evolution on *Chlamydomonas reinhardtii* traits. We replicated each gradient level 4 times, with the exception of salt gradient levels, in which each population was replicated only once.

| Level | Nitrogen gradient (μM N) | Phosphorus gradient (μM P) | Light gradient (μM photons $m^{-2}s^{-1}$) | Salt gradient (g/L NaCl) | Thermal gradient (°C) |
|---|---|---|---|---|---|
| 1 | 5 | 0.5 | 0.25 | 0 | 10 |
| 2 | 10 | 1 | 1.5 | 1 | 16 |
| 3 | 20 | 2 | 5 | 2 | 22 |
| 4 | 40 | 4 | 12.5 | 3 | 28 |
| 5 | 60 | 6 | 27.5 | 4 | 34 |
| 6 | 80 | 8 | 50 | 5 | 40 |
| 7 | 100 | 10 | 82.5 | 6 | |
| 8 | 400 | 20 | 125 | 7 | |
| 9 | 600 | 35 | 175 | 8 | |
| 10 | 1000 | 50 | 250 | 9 | |

**Table S9. Information on model fits for different thermal performance curves.**

We used Akaike Information Criterion values to select among a range of commonly used phenomenological and mechanistic thermal performance curve (TPC) models (7). For comparative purposes, we fit all models to every experimentally evolved and ancestral *C. reinhardtii* population using the rTPC package (6) and report the mean value for each model. We selected the Lactin II model, which fell within two AIC units of the best fitting model (ΔAIC < 2) and, unlike the marginally better scoring models, has the capacity to model negative growth rates at extreme temperatures.

| Model name | Mean AIC score |
|---|---|
| Analytis-Kontodimas | 32.18 |
| Ashrafi II | 66.58 |
| Atkin | 32.18 |
| Briere | -0.46 |
| Deutsch | -3.18 |
| Eubank | 49.68 |
| Lactin II | -1.97 |
| Mitchell-Angilletta | 53.20 |
| Ratkowsky (bounded) | -2.96 |
| Rezende | -1.13 |
| Taylor-Sexton | 25.63 |
| Thomas I | 1593.27 |

**Table S10. Complete list of R2jags model priors.**

We fit Bayesian models in R2jags (5) to estimate variation in the exponential growth rates of *Chlamydomonas reinhardtii* across light, nitrogen, phosphorus, temperature, and salinity gradients and extract key niche-determining trait values.

| Gradient | Model | | | | |
|---|---|---|---|---|---|
| Light | Monod curve | $\mu_{max}$ : [0 – 5] | $k_s$ : [0 – 10] | | |
| Nitrogen | Monod curve | $\mu_{max}$ : [0 – 5] | $k_s$ : [0 – 10] | | |
| Phosphorus | Monod curve | $\mu_{max}$ : [0 – 5] | $k_s$ : [0 – 10] | | |
| Temperature | Lactin II TPC | ρ : [0 – 0.2] | $T_{max}$ : [36 – 44] | $\Delta T$ : [0.1 – 6] | λ : [-3 – 0] |
| Salinity | Inverse logistic growth | *a* : [0.5 – 2] | *b* : [0 – 10] | *S* : [0 – 10] | |
